# The PVD neuron has male-specific structure and mating function in *C. elegans*

**DOI:** 10.1101/2024.05.19.594847

**Authors:** Yael Iosilevskii, David H. Hall, Menachem Katz, Benjamin Podbilewicz

## Abstract

Neurons display unique shapes and establish intricate networks, which may differ between sexes. In complex organisms, studying sex differences in structure and function of individual neurons is difficult. The nematode *Caenorhabditis elegans* hermaphrodites and males present an exceptional model for studying neuronal morphogenesis in a simple, sexually-dimorphic system. We focus on the polymodal sensory bilateral neuron pair PVD, which forms a complex but stereotypic dendritic tree composed of multiple subunits that resemble candelabra. PVD is well studied in hermaphrodites, but not in males. We show here that during larval development, male PVDs extend a similar architecture to the hermaphrodite utilizing the sexually-shared Menorin patterning mechanism. In early adulthood, however, male PVD develops a unique extension into the copulatory tail structure. Alongside established tail ray neurons RnA and RnB, we show PVD is a third, previously unrecognized, neuron within the tail rays. Unlike RnA and RnB, PVD extends anterogradely, branches and turns within the ray hypodermis, and is non-ciliated. This PVD sexually-dimorphic arborization is absent in mutant backgrounds which perturb the Menorin guidance complex. SAX-7/L1CAM, a hypodermal component of this complex, shows a male-specific expression pattern which precedes PVD extension, and its presence allows PVD to enter the tail rays. Further, our results reveal that genetically altered arborization or ablation of the PVD result in male mating behavioral defects, particularly as males turn around the hermaphrodite. These results uncover an adult-stage sexual dimorphism of dendritic branching and uncover a function for PVD in male sexual behavior.

**Significance Statement:** Neurons form intricate shapes and networks, which may display sexual differences. Pinpointing these changes at the single cell level, however, is challenging. In *C. elegans*, the PVD neuron is a powerful model for stereotypical neuron shaping, yet most studies concern only one of two sexes. Our research focuses on the understudied male, where we show an adult-stage extension of PVD into the male copulatory tail organ. We further show how this stems from a sex-shared patterning complex, progressing independently and in a different environment compared with nearby male-specific neurons. We further find PVD has a role in the male adult-specific mating behavior. PVD thus presents a unique example of a highly arborized neuron showing sexually dimorphic behavioral function and structure.

## Introduction

Neurons grow distinctive morphologies and form sophisticated networks (1–3). These networks often show sexual dimorphism, with different connectivity and distribution between the sexes (3–6). In humans, multiple neurological disorders, including Parkinson’s disease and dementia, are more prevalent in one sex (7). While altered neuron shape, in particular dendritic spines, has been associated with some of these disorders (8), these differences are difficult to attribute to any sex-specific dendrite morphology.

The nematode *Caenorhabditis elegans* has been widely used as a model organism for studying neuron shaping (2, 3), connectivity (2, 9), plasticity (10–13) and activity (9, 14). While the majority of the wild type population are hermaphrodites, loss of one X chromosome yields males (15), which are sexually dimorphic in anatomy (16), neuronal connectivity (3), and behavior (15). Both the hermaphrodite and the male’s compact nervous system consists of diverse neuron types and morphologies, which enable a variety of behaviors (5), including oriented movement (17) and escape (9, 18, 19). Uniquely, each neuron’s identity, and the connectome linking them together, are fully established (2, 3). Of these, a small subset of neurons in *C. elegans* present highly arborized dendrites, providing a robust system for analyzing reproducible neuron patterning and development (20), among them the neuron pair PVD (21–23).

In hermaphrodites, each of two PVD neurons (left and right) assumes its structure during the final three out of four larval developmental stages (L2-L4) (21, 22) (Fig. 1 *A-C*). PVD is a polymodal sensory neuron, responding to noxious touch (18, 21), temperature drops (24), sound vibration (19) and body position (25, 26). Its characteristic arborization, forming repetitive subunits which resemble candelabra (‘menorahs’ (21)), develops through actin-mediated growth (27). The hermaphrodite PVD morphogenesis is guided by the Menorin multipartite complex, in which transmembrane receptors from the PVD, notably the leucine-rich repeat transmembrane domain (LRR-TM) protein DMA-1 (28), bind hypodermal (epidermal) and muscle-secreted guidance cues, notably SAX-7/L1CAM, MNR-1/Menorin and LECT-2/Chondromodulin II (27, 29, 30). Its initial branching is governed in part by a LIM homeobox domain transcription factor, MEC-3 (22, 31–33). In addition to the Menorin patterning complex, multiple pathways maintain PVD tiling, self-avoidance, and prune excessive branching events, among them UNC-6/Netrin (34, 35), KPC-1/Furin (36) and the cell-cell fusogen EFF-1 (21, 37). With age and following dendritic injury, PVD loses its characteristic ordered form and develops additional ‘ectopic’ branches, which extend beyond the young-adult menorah shapes (12, 38) (Fig. 1 *C*).

**Figure 1.**
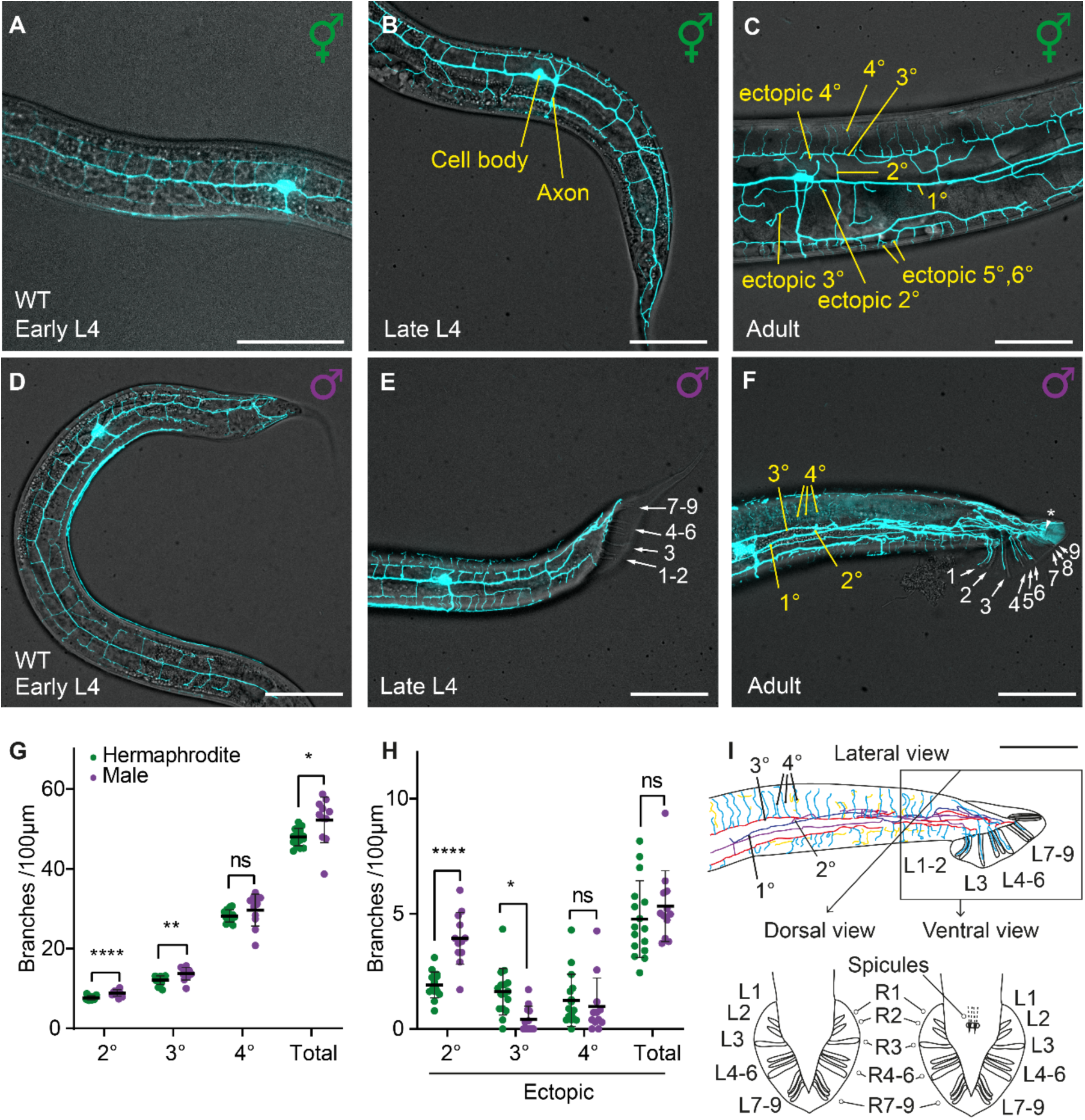
WT male PVD development shows post-larval sexual dimorphism. (*A-C*) Hermaphrodite PVD development in L4 through to adult. PVD branch orders are marked yellow in panel *C*. Anterior is left, ventral down. *ser2prom3::GFP* PVD marker. Scale bar is 50 µm. (*D-F*) Male PVD development in L4 through to adult; white numbers and arrows note tail rays; PVD branch orders are marked in yellow in panel *F*. *ser2prom3::GFP* PVD marker. Asterisk notes PVD branching into the phasmids. Scale bar is 50 µm. (G) Analysis of PVD branches. *t* test, Holm-Šídák multiple comparison correction. Black bars show mean, error bars show ± SD. *P* <0.05 (*), *P* <0.01 (**), *P* <0.001 (***); *P* <0.0001 (****). (H) Ectopic branches around the cell body region in 1-day adult animals, normalized per 100 µm of the primary branch for the different branching orders; n = 16 (6 PVDL,10 PVDR) hermaphrodite and n = 12 (7 PVDL, 5 PVDR) males. *t* test, Holm-Šídák multiple comparison correction. Black bars show mean, error bars show ± SD (except male ectopic tertiaries and ectopic quaternaries, showing + SD). (I) Cartoon depiction of the male tail organ; lateral view contains a tracing of the neuron in panel *F*: projection of three-dimensional trace, branch orders annotated blue (1°), purple (2°), red (3°), cyan (4°) and yellow (5° and 6° order ectopics); scale bar is 50 µm; PVD branch orders are marked in black following panel *F*. Ventral and dorsal cartoons are not drawn to scale.

Sexually dimorphic behavior in *C. elegans*, typically unique to the male, can arise from altered expression of sensory channels (39, 40), variation in synaptic connectivity and strength (41, 42) and different gene expression networks (43, 44), some of which may already be active prior to sexual maturation (45). To date, differences in dendritic arborization between hermaphrodite and male *C. elegans* have been characterized in the motor neuron PDB (46), but, to our knowledge, never in a sensory neuron. Here, we focus on PVD, and ask whether its complex arbor has any sexually dimorphic features. We show PVD develops a post-larval, male-specific elaboration of its dendrite morphology, which enters into the specialized rays of the male tail, previously thought to contain only two neurons (RnA and RnB) (16). Our results reveal that, unlike the ciliated RnA and RnB, PVD is non-ciliated, extends anterogradely, and branches within the ray hypodermis. Its male-specific post-larval morphogenesis is consistent with a sexually dimorphic distribution of the hypodermal patterning cue SAX-7/L1CAM, acting within the sexually-shared Menorin patterning complex (28–30, 36). We further show that in addition to its sensory functions in hermaphrodites, PVD is required for precise male mating behavior. Our results uncover sexual dimorphism in PVD morphogenesis and function, which also open directions for studying adult-onset directed dendritic branching.

## Results

### PVD morphology is sexually dimorphic

While much is known about hermaphrodite PVD dendritic tree morphogenesis (Fig. 1 *A-C*) (21, 22), it remains unclear whether males share the same PVD arborized structure, patterning mechanism, and function. To determine the wildtype (WT) structure of the male PVD, we imaged animals either from stocks continuously maintained with males or carrying a *him-5(e1490)* background mutation which increases the male percentage (15). We found that male PVD develops candelabra-shaped dendritic units during the latter larval stages, L3 and L4, similar to the hermaphrodite (Fig. 1 *A-F*). Four branching orders are maintained; the male primary branch is compressed medially compared with the hermaphrodite (Movies S1, S2). Branch density relative to body length is higher in males (Fig. 1 *G*), with altered distribution of additional ‘ectopic’ branching (Fig. 1 *H*). Interestingly, we noted that in adult males, PVD extends further and enters into the sexually dimorphic tail rays (Fig. 1 *F*), confirmed by flow of photoconvertible Kaede protein from the cell body (Fig. S1). The nine bilateral male tail rays support a specialized fan (Fig. 1 *I*) which is essential for mating (47, 48). Each ray has been shown to contain only two unique neurons, RnA and RnB (16, 49), prompting us to investigate this novel placement of the PVD further. We found that PVD does not enter the rays as they first form in the fourth larval stage (L4); rather, PVD ray extension is initiated around the final L4-to-adult molt (Fig. 1 *E* and *F*). To determine whether this branching phenomenon is intrinsic to the male identity of PVD itself, we utilized cell-specific feminization (50) by expressing a dominantly active intracellular domain of the TRA-2 receptor from a PVD promoter. This analysis revealed that the male-specific branching into the tail rays is not dependent on the sex of PVD itself (Fig. S2), leading us to look into the effect of the surrounding cells on its morphogenesis.

### PVD dendrite entry into tail rays is affected by tail morphology and the fusogen EFF-1

Each ray of the male tail contains a hypodermal component, a glia-like structural cell, and two specialized ray neurons (16, 51). To study whether the structural cell directs the PVD entry into the rays we sought to perturb the ray-forming cell sublineage, by using *lin-44(n1792)* Wnt ligand mutants. In these mutants the structural cell is likely misspecified, and the final number of rays is reduced (52). We found that defects in this morphogen partially hindered PVD entry, yet did not abrogate it (Fig. S3), suggesting the structural cell alone may not be strictly necessary for PVD processes to progress into the tail rays.

To determine how overall tail morphology and PVD dendrite extension interact, we next examined mutants of the epithelial fusogen *eff-1* which governs the correct formation of the syncytial hypodermis (53). Male *eff-1* mutants also show deformed tails, with a small fan and fewer, shorter rays (53). In hermaphrodite *eff-1* mutants PVD is uniquely hyperbranched, extending multiple ectopic processes (21), a phenotype we noted is also true for males (Fig. S4 *A-C*). This led us to speculate that, despite the hypodermal defects, this excessive branching may translate to increased, arbitrary, ray entry. In males, however, this is not the case (Fig. S4). Rather, loss of *eff-1* inhibits PVD ray extension in a dosage-dependent manner, as established by comparison of *eff-1(hy21ts)* temperature-sensitive allele at restrictive and permissive temperatures (Fig. S4 *A-D*). By analyzing several ages, it appears PVD branches within the rays are still dynamic and growing (Fig. S4 *E*), indicating the effect likely stems from the surrounding ray structure. We conclude that PVD entry into the tail rays is influenced by the overall tail morphology, likely depending on the correct formation of the hypodermal ray component and possibly the structural cell as well.

### PVD intra-ray branching is independent of RnB ray neurons

To date, two neurons innervating the male tail rays were identified, named RnA and RnB (each numbered n =1-9 and bilateral; R1A(R/L), R2A(R/L) etc.) (16). To study whether ray ‘resident’ neurons such as RnBs and PVD branches interact during tail morphogenesis, we constructed strains co-expressing distinct fluorescent reporter proteins for RnB and PVD. We found that in WT L4 animals, as the rays are spun from the body while the tail is retracting (51), RnB is already present within each ray, while PVD is absent (Fig. 2 *A*, n = 54 rays, Movie S3). PVD extends processes into the rays only in adults, where we note PVD and RnB signals remain spatially separated (Fig. 2 *B* and *C,* Movie S4). This is also true for *lin-44(n1792)* mutants (Fig. S3 *C* and *D*). In order to further determine whether RnB presence is strictly required for subsequent PVD entry, we utilized a strain where RnB neurons are killed by specific ICE human caspase expression (54). We find that in these animals PVD retains the ability to enter the tail rays regardless of RnB presence (Fig. S5). We conclude that RnB and PVD morphogenesis are separated both temporally and spatially, revealing differential mechanisms of intra-ray dendritic growth: while RnB uses a retrograde extension during the L4, PVD uses an anterograde extension upon adulthood.

**Figure 2.**
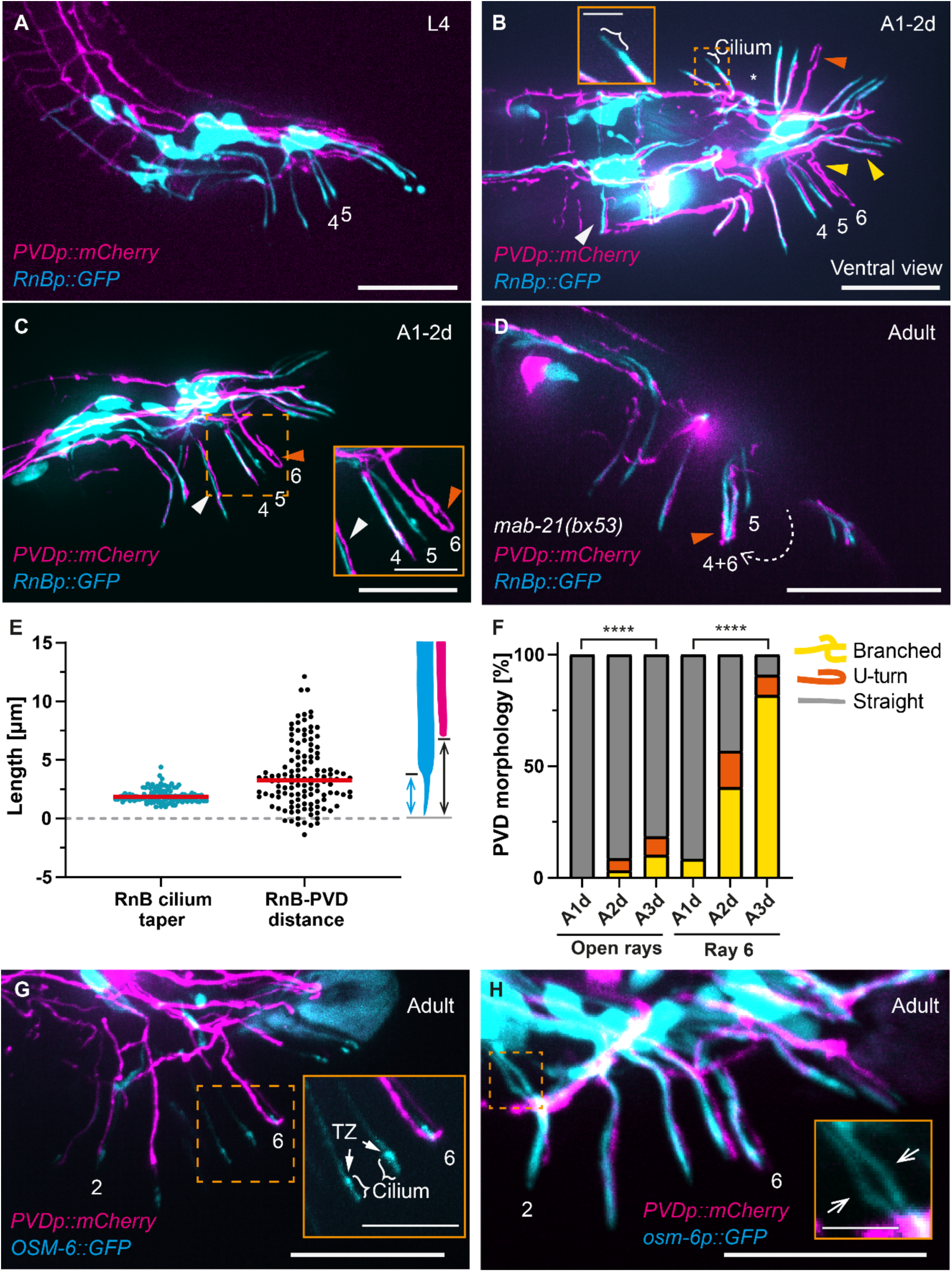
PVD processes are not ciliated and enter the rays after, but not over, RnB ciliated ray neurons. (*A-D*) Examples of *him-5(e1490)* male tails expressing the RnB neuron marker *pkd-2::GFP* (expressed in all but R6B; cyan), and *pF49H12.4::mCherry* PVD marker (magenta). (A) Lateral view of an L4 WT. RnB dendrites enter the rays using a retrograde mechanism (23, 51, 72) before the PVD initiates its anterograde extension. Rays 4, 5 marked for orientation purposes. Scale bar is 25 µm. (B) Ventral view of an adult WT (1-2 day, A1-2d). In the majority of rays and RnB axon paths (62), PVD (*pF49H12.4::mCherry*) and RnB (*pkd-2::GFP*) processes show zero overlap (75/144 RnB dendrites and 65/66 RnB axons), with partial-length, minimal overlap in the remaining cases. When quantifying tapered tip morphology in these animals, 3/67 ray PVD processes show thinner endings compared with 101/124 RnB dendrites. A ciliated tip example of R2B (top) is highlighted by a white bracket in both the panel and its inset. Rays 4, 5, 6 indicated for orientation. Orange arrowhead highlights a U-turn example, yellow arrowheads highlight intra-ray branching, white arrowhead points to a non-overlapping PVD dendritic branch and R2B/R3B axon(s) (62). One ray 3 is missing (occasional in WT, (16)) with R3B localized to its base and PVD projecting alongside (asterisk, top). Scale bar is 25 µm. Inset scale bar is 5 µm. (C) Lateral view of an 1-2 day adult (A1-2d) WT *him-5(e1490)* animal expressing PVD (*pF49H12.4::mCherry*) and RnB (*pkd-2::GFP*) markers. Rays 4, 5, 6 are indicated for orientation. Orange arrowhead highlights a U-turn example, white arrowhead points to a non-overlapping PVD dendritic branch and R3B dendrite, in both the panel and its inset. Scale bar is 25 µm. Inset scale bar is 10 µm. (D) Lateral view of an adult *mab-21(bx53); him-5(e1490)* male expressing PVD (*pF49H12.4::mCherry*) and RnB (*pkd-2::GFP*) markers. Rays 4 and 6 are ‘fused’, with ray 6 translocating anteriorly to ray 5, to join ray 4 (dashed arrow). Note that, unlike the WT (62), this fusion causes ray 6 to express *pkd-2::GFP* (ray 6 *pkd-2::GFP* signal appears in 15/15 *mab-21(bx53)*, 0/23 WT). A single PVD branch enters the fused ray and performs a U-turn (orange arrowhead). Scale bar is 25 µm. (E) Approximate RnB cilium lengths (cyan) and distance from the distal end of PVD to the tip of RnB (black) in WT *him-5(e1490)* 1-2 day adults expressing *pkd-2::GFP* RnB marker (cyan cartoon) and *pF49H12.4::mCherry* PVD marker (magenta cartoon), measured based on fluorescent signal. R6B is not analyzed owing to absence of *pkd-2::GFP* signal. n = 116 tapering RnB measurements and 127 RnB-PVD dendrite pair distances. Red lines indicate medians. (F) Progression of intra-ray branching (yellow) and U-turn (orange) phenotypes of PVD within open rays (*i.e.* all but ray 6, left), and the closed ray 6 (right), during the first three days of adulthood, in WT *him-5(e1490); ser2prom3::GFP* males. A U-turn is defined by the distal tip executing a complete folding back and continuing medially towards the body, whereas a Branched process is any forked endpoint extending from a branch within the ray. A1d = one day adult; A2d = two day adult; A3d = three day adult. n = 62,186,108 open rays and n = 12, 37, 22 ray 6 cases scored for the three age groups, respectively. Fisher’s exact test comparing A1d to A3d and straight to ‘wandering’ branched, (****) *P* <0.0001. (F) PVD does not express the ciliated neuron reporter *osm-6p::OSM-6::GFP*. Tail region of an adult *him-5(e1490)* male expressing *osm-6p::OSM-6::GFP*, targeted to ciliated endings, and *pF49H12.4::mCherry* PVD marker. No colocalization was detected in n = 94 rays observed across 13 animals. Rays 2 and 6 indicated for orientation. Scale bar is 25 µm. Inset of dashed orange rectangle: examples of two cilia (brackets) and their transition zones (TZ). Scale bar is 10 µm. (G) Tail region of an adult *him-5(e1490)* male expressing *osm-6p::GFP*, expressed in ciliated neurons including PDE and PQR, and *pF49H12.4::mCherry*, expressed in PVD, AQR and PQR. 0/15 adult males show colocalization of the two markers in the PVD soma (see also Fig. S6). Rays 2 and 6 indicated for orientation. Scale bar is 25 µm. Inset of dashed orange rectangle: example of R2A and R2B converging in a ray. White arrows in the inset point to the two neurons. Scale bar is 5 µm.

### PVD dendrites in the rays are not ciliated

RnA and RnB are both ciliated, and project in a neuronal channel formed by the structural cell, opening to the external surrounding in all rays but 6 (16, 55). The presence of PVD alongside these neurons led us to wonder whether PVD similarly extends ciliated endings and opens to the environment. In following PVD extension into the rays, we noticed PVD processes further branching within each ray, and, at times, performing a U-turn before reaching the ray tip (Figs. 2 *B, C* and *F*). PVD often does not extend to the very tip of the ray (Fig. 2 *E*) and does not taper into the recognizable cilium shape readily observed in RnB (Fig 2 *B*). Moreover, PVD fails to express the ciliated neuron reporter OSM-6 nor the transcriptional reporter *osm-6p::GFP* (56, 57) in both hermaphrodites and males (Figs. 2 *G* and *H* and S6). Based on recent neuronal cell-specific RNA sequencing analyses, cilia-related RNAs, including those encoding structural components (*tax-2/*cyclic nucleotide gated olfactory channel), motor components (*che-3/*dynein, *osm-3*/kinesin-II and *xbx-1*/dynein light intermediate chain) and intraflagellar transport-related genes (*bbs-1*, *bbs-2*, *bbs-5*, *bbs-8*, *che-2*, *che-11*, *che-13*, *ift-81*, *osm-5*, and *osm-6*) cannot be detected in PVDs of both males and hermaphrodites (58, 59) (*Appendix SI* Table S4). Based on these results we conclude that male PVD processes in the rays remain non-ciliated, and do not open to the external environment.

### PVD can occupy the full width of a ray

Having noted that PVD may U-turn and branch within rays, we next sought to characterize this morphology and determine which cells PVD may interact with within the ray. Our analysis of different ages reveals that this intra-ray arborization is enhanced with time, and that PVD may U-turn in rays which are otherwise open to the surrounding (Fig. 2 *F*), further strengthening our assumption that PVD processes are not directed towards an external pore.

The age-dependent post-larval extension of PVD into the rays (Fig. 2 *A* and *B*) presents a unique model to explore dendritic sculpting. To determine the effect of ray placement on this phenomenon, we first studied *mab-20(bx61ts)* mutants, which present an increased percentage of joined (‘fused’) rays (60), where two or more rays are displaced and join to share an epidermal sheath (49). Our analysis of these joined rays revealed they often contained a single PVD dendritic process, which subsequently branches or U-turns (Fig. S7). In the WT, ray 6 is naturally wider compared with the rest (60), and shows the same phenomenon (Fig. S7 *B*), which can be followed by live imaging (Fig. S8). This suggests ray placement has little effect on subsequent PVD extension, yet wider rays allow a single PVD dendrite to branch and U-turn further (Figs. S7 and S8).

In order to further probe which cells may interact with PVD during ray extension, we utilized a second fused-ray mutant, *mab-21(bx53)*, where electron microscopy studies have validated the presence of two channels within a fusion of ray 6 into ray 4. These two channels are surrounded by two fused structural cells, and contain two separate RnBs and one or two RnAs (61). We hypothesized that if PVD processes are instructed from the channels, then we would observe two PVD processes in the fused rays. Unlike the WT (62), the displaced *mab-21(bx53)* ray 6 surprisingly expresses the RnB *pkd-2::GFP* marker (Fig. 2 *D*), allowing us to determine whether PVD interacts with both neuron channels within a single ray. We found that in those cases, PVD tends to send a single process (7/11 cases) rather than two, and remains flexible within the ray width (Fig. 2 *D*). Thus, it is unlikely that PVD processes require interaction with the ray’s channel(s). Combining our results from two fused-ray mutant backgrounds and the different ray width variability of the WT (Figs. 2 and S7), together with PVD-RnB fluorescent marker spatial separation (Movie S4), we suggest that PVD extends outside of the ray neuronal channel.

### Ray ultrastructure reveals PVD processes are outside the channel

To directly establish which cells physically interact with intra-ray PVD dendrites, we turned to previously unpublished electron microscopy cross sections of a young male adult WT ray 6, and were able to locate a third neuronal process, distinct from RnA and RnB, progressing between the structural cell and the hypodermis (Fig. 3 *A* and *B*). This provides direct evidence for the existence of three neurites within a wild-type male tail ray. To further determine whether PVD has any ciliated neuron ultrastructural characteristics, we performed EM analysis of hundreds of ray sections from multiple young adult male tails. These revealed that when seen, male cilia in open rays are always at the ray tip, one embedded in the structural cell, and the second always emerging out the pore. Notably, in every case observed there are exactly two ciliated endings, with only one finding room to fit into the pore (Fig. S9 *A* and *C*). We propose that if PVD regularly formed a third cilium within most rays, such an example would have been noted (16, 61, 63). The PVD dendrite morphology we observed shares some features with PVD quaternary branches elsewhere (21), although ray processes vary in their width, and may be twice the diameter of a typical PVD quaternary branch (Figs. 3 *B* and S9 *B*) or even thinner than one (Fig. S9 *C*). In addition to its clear cytoplasm, which is distinctively emptier than the rays and without mitochondria or vesicles, we note PVD shows no microtubules (Fig. S9), further suggesting it would not form a ciliated ending. We conclude PVD processes enter the male tail rays outside the neuronal channel (Fig. 3 *A* and *B*), and remain within the ray hypodermal component without opening to the external environment.

**Figure 3.**
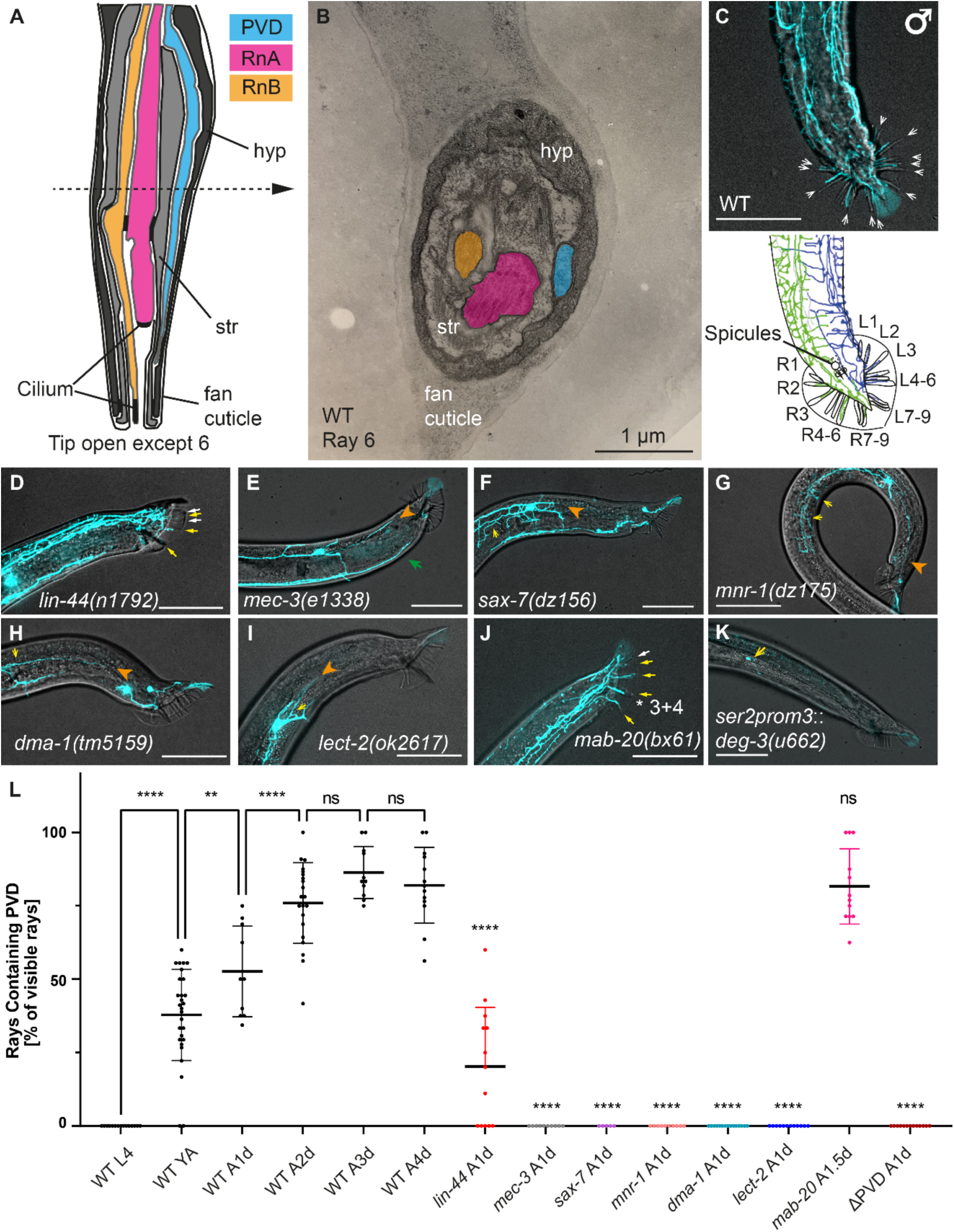
Male PVD enters the tail rays outside the neuron channel and shares mutant phenotypes with hermaphrodite-characterized morphogenesis genes. (A) Cartoon depiction of a longitudinal section across an open male tail ray, noting hypodermis (hyp), structural cell (str), fan cuticle, ray neurons RnA (magenta), RnB (orange) and their cilia, and PVD (cyan). Based on (16). (B) Transmission electron microscopy image of a thin cross section of a young adult WT ray 6, noting hypodermis (hyp), structural cell (str), fan cuticle and ray neurons RnA (magenta), RnB (orange) and PVD (cyan) as in panel *A*. Scale bar is 1 µm. (C) Top: ventral view of a two-day adult male, *ser2prom3::GFP* PVD marker. Scale bar is 50 µm. Bottom: cartoon tracing of respective PVDL (blue) and PVDR (green). Arrows denote PVD branching into the rays. (D) Male one-day adult *lin-44(n1792); him-5(e1490)* with *ser2prom3::GFP* PVD marker. Yellow arrows indicate a ray with PVD signal, white arrows indicate rays without PVD signal. Scale bar is 50 µm. (E) Male one-day adult *mec-3(e1338); him-5(e1490)* expressing *ser2prom3::GFP* PVD marker, orange arrowhead marks the posterior-most branch. A green arrow points to a process of the PDE (31, 77), most visible here but also present in panel *D* and Fig. 1 *D* and *E* (also see Materials and Methods). Scale bar is 50 µm. (F) Male one-day adult *sax-7(dz156)* expressing *pF49H12.4::GFP* PVD marker, yellow arrows point to examples of characteristic altered structure, orange arrowhead marks the posterior-most branch. Scale bar is 50 µm. (G) Male one-day adult *mnr-1(dz175)* expressing *pF49H12.4::GFP* PVD marker, yellow arrows point to examples of characteristic altered structure, orange arrowhead marks the posterior-most branch. Scale bar is 50 µm. (H) Male one-day adult *dma-1(tm5159)* expressing *pF49H12.4::GFP* PVD marker, yellow arrow points to characteristic altered structure, orange arrowhead marks the posterior-most branch. Scale bar is 50 µm. (I) Male one-day adult *lect-2(ok2617)* expressing *ser2prom3::GFP* PVD marker, yellow arrow points to characteristic altered structure, orange arrowhead marks the posterior-most branch. Scale bar is 50 µm. (J) *mab-20(bx61ts); him-5(e1490) 1-2-day adult* expressing *ser2prom3::GFP* PVD marker at a semi-restrictive 22°C temperature, asterisk marks a characteristic ‘fusion’ joining rays 3 and 4. Yellow arrows indicate rays with PVD signal, white arrows indicate rays without PVD signal. Scale bar is 50 µm. (K) PVD genetic ablation *ser2prom3::deg-1(u662); him-5(e1490)* two-day adult expressing *ser2prom3::Kaede* PVD marker showing signal remnant (arrow) next to the PVD soma, corresponding to PDE, with ventral truncated axon-like process. Scale bar is 50 µm. (L) Quantification of PVD entry into the tail rays in different mutant backgrounds and ages; all animals assayed express the integrated *ser2prom3*::*GFP* PVD marker and are *him-5(e1490*), with the exception of *deg-3(u662)* genetic ablation which utilizes a Kaede fluorescent marker, *lect-2(ok2617)* which has no *him-5* background, and *sax-7(dz156), mnr-1(dz175)* and *dma-1(tm5159)* which express an integrated *pF49H12.4::GFP* PVD marker and are without *him-5(e1490)*; YA: young adult; A1d, A1.5d, A2d, A3d, A4d: 1, 1.5, 2, 3 and 4-day adult, respectively. Day 1 adult = L4 + 1 day. n = 15, 28, 10, 21, 11, 13, 13, 10, 5, 10, 15, 12, 12, 13, respectively. For *mab-20(bx61ts)*, one day from L4 at room temperature is comparable with to L4 + 28 h at 20°C and as such marked 1.5 day adult. One-way ANOVA, Šídák multiple comparison correction. Black bars show mean, error bars show ± SD. *P* <0.05 (*), *P* <0.01 (**), *P* <0.001 (***); *P* <0.0001 (****).

### Hermaphrodite-characterized PVD patterning mutants show similar male phenotypes

The hermaphrodite PVD morphogenesis is guided by the Menorin multipartite complex, in which transmembrane receptors from the PVD, notably the leucine-rich repeat transmembrane domain

(LRR-TM) protein DMA-1 (28), bind hypodermal (epidermal) and muscle-expressed guidance cues, notably SAX-7/L1CAM and MNR-1/Menorin, alongside LECT-2/Chondromodulin II (27, 29, 30). Its initial branching is governed in part by a LIM homeobox domain transcription factor, MEC-3 (33). To characterize whether male PVD retains these properties within the tail fan region, we first imaged a ventral view of a WT adult tail (Fig. 3 *C*, top) and traced both left and right PVD neurons (Fig. 3 *C*, bottom). This revealed male PVDs maintain lateral tiling as in the hermaphrodite (21, 64), extending into their respective sides (Fig. 3 *C*). Similarly to the progression noted for *eff-1* mutants (Fig. S4 *E*), WT male PVD gradually enters the tail rays in an age dependent fashion, beginning from the newly molted young adult (Fig. 3 *L*).

To determine whether its patterning mechanism is sexually shared, we imaged *mec-3(e1338), sax-7(dz156), mnr-1(dz175), dma-1(tm5159)* and *lect-2(ok2617)* mutant male animals and found that these backgrounds show similar PVD morphogenesis defects to hermaphrodites (22, 29, 30, 33) (Fig. 3 *E-I*). We also confirmed PVD genetic ablation has a similar effect in males (Fig. 3 *K*) (25). These results reveal that the roles of *mec-3* and the Menorin guidance pathway in PVD morphogenesis are sex-shared. We note that in these mutant backgrounds PVD also fails to extend into the tail rays (Fig. 3 *L*), in contrast to *lin-44(n1792)* and *mab-20(bx61ts)* (Fig. 3 *D* and *J* and *L*).

### SAX-7 hypodermal patterning guides PVD dendrite entry into the male tail rays

Noting the arborization defects seen in Menorin complex mutant backgrounds also inhibit PVD extension into the tail rays (Fig, 3 *L*), we wondered whether its patterning components are present in the developing rays, notably the axonal guidance molecule SAX-7/L1CAM (23, 29, 30). We imaged males carrying an integrated low-copy number fosmid-based reporter for *sax-7* translation (65) and found that during L4, a *sax-7p*::*sax-7::GFP* signal is visible within the nascent rays (Fig. 4 *A-D*). This patterning precedes PVD extension (Figs. 3 *L*). In the adult, PVD signal follows over the tail ray SAX-7::GFP pattern, which is prominent in all the rays (as also shown by (66)), (Fig. 4 *C-E*, n =113 rays). In addition to the loss of PVD branching in the rays of *sax-7* mutant animals (Fig. 4 *F*), this branching was restored by hypodermal-specific expression of the short SAX-7 isoform (Fig. 4 *F-H*), previously used to rescue the hermaphrodite PVD morphogenesis defects of *sax-7* mutants (29, 30). Thus, in male adults, *sax-7* expression in the ray hypodermis appears to facilitate the entry of PVD branches into the tail. These results suggest a model whereby the sexually dimorphic aspect of PVD morphogenesis utilizes components of the established intercellular guidance complex in the hermaphrodite (23, 29, 30), however with a male-specific distribution and at a heterochronic post-larval stage.

**Figure 4.**
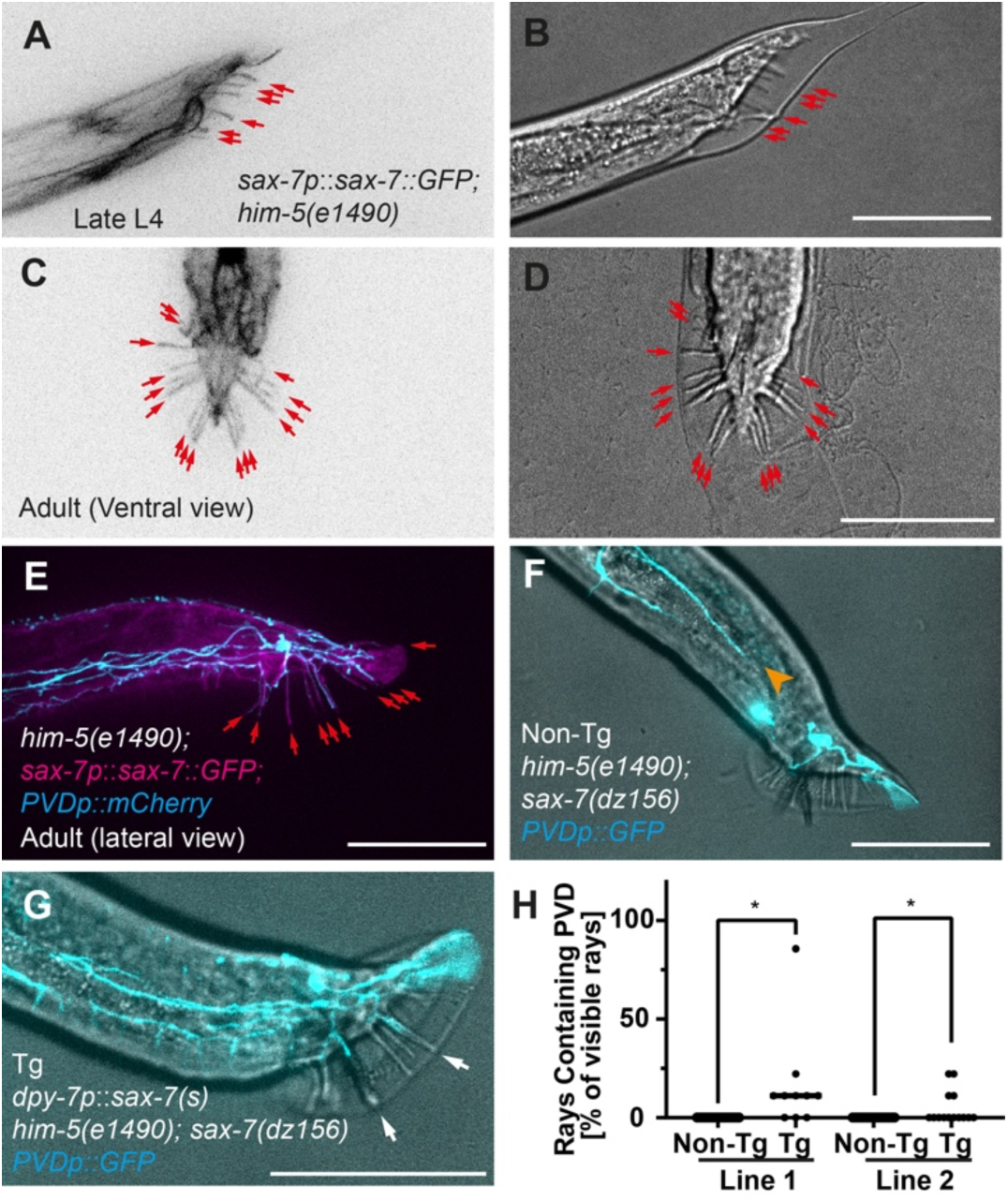
SAX-7 in the hypodermis is required for PVD extension into the rays. (*A,C*) Male tail with *sax-7p::sax-7::GFP* reporter (inverted) in L4 (*A*) and adult (*C*). Signal observed in 43/43 nascent (L4) rays. *(B,D*) Matching DIC images to A and C, respectively. Red arrows point to tail rays. Scale bar is 50 µm. (*E*) Adult lateral view of merged *sax-7* reporter (magenta) with PVD marker (cyan), ‘inverted’ pseudo-colors are used for clarity. 113/113 one-day adult rays observed with SAX-7 signal, of which 79/113 expressed PVD signal. (*F*) One day adult *sax-7(dz156); him-5(e1490)* male expressing *pF49H12.4::GFP* PVD marker, non-transgenic (Non-Tg) control for animals carrying a *dpy-7p::sax-7(s)* short variant transgene (29, 30). Orange arrowhead marks the posterior-most branch, non-transgenic (Non-Tg) control. Scale bar is 50 µm. (*G*) One day adult *sax-7(dz156); him-5(e1490)* male expressing *pF49H12.4::GFP* PVD marker, transgenic (Tg) animal expressing *dpy-7p::sax-7(s)* short variant (29, 30), determined by co-injection marker (pharyngeal muscle *myo-2p::GFP*) presence. White arrows show PVD entry into rays 3 and 6. Scale bar is 50 µm. (*H*) Analysis of PVD entry into the male tail rays in animals carrying extrachromosomal *dpy-7p::sax-7(s)* short variant (29, 30) comparing transgenic (Tg) with non-transgenic (Non-Tg) animals based on co-injection marker expression (pharyngeal muscle *myo-2p::GFP*). Unpaired *t* test, *P* < 0.05 (*). A1d one-day adult animals are analyzed from two independent transgenic lines (n = 16, 10, 16, 14 respectively).

### PVD initiates dendritic branching into the rays prior to the onset of male mating behavior

Male mating in *C. elegans* is a complex multi-step process, performed by a male animal upon encountering a hermaphrodite (48). Its execution relies on male-specific neurons and neural circuits (5) which guide chemosensory sexual attraction (39) and mechanosensation (48, 67, 68) as well as on sexually-dimorphic anatomy, in particular the male tail (16, 48, 68). Once it has located a hermaphrodite, the male circles around it tail-first until the tail makes contact with the hermaphrodite’s vulva (48). The male then inserts its two copulatory spicules and ejaculates sperm into the hermaphrodite’s uterus (48). Our results suggest PVD is neither ciliated nor open, and unlikely to be chemosensory, leading us to focus on steps in male mating behavior which may involve mechanosensation (18, 21, 32) and proprioception (25, 26). We chose to analyze three behavioral steps: 1) contact recognition, which is the initial reversal in response to tail contact with a hermaphrodite; 2) precise turning around the head or tail of the hermaphrodite during backwards movement to maintain scanning for the vulva; and 3) stopping at the vulva (Fig. 5 *A*) (48).

**Figure 5.**
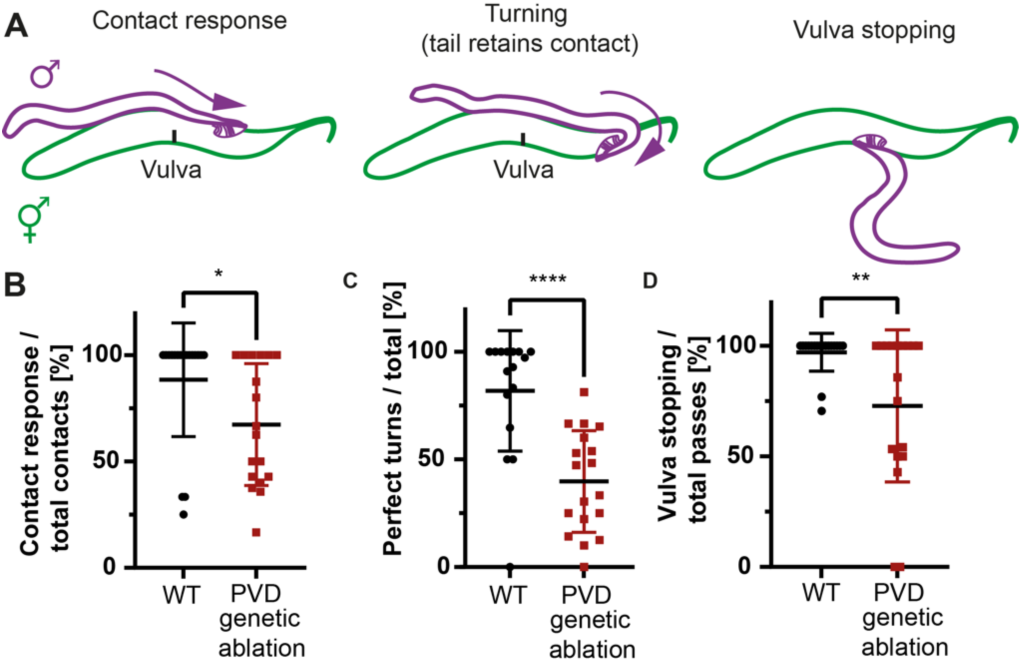
Mating behavior is impaired under PVD genetic ablation. (A) Cartoon of mating steps. Male presented in purple, hermaphrodite in green. (B-D) Hermaphrodite recognition, turning and vulva recognition, respectively, in WT and *ser2prom3*::*deg-3(u662)* animals where PVD undergoes early degeneration. B: n = 18 WT, 18 *ser2prom3*::*deg-3(u662)*; C: n = 16 WT, 18 *ser2prom3*::*deg-3(u662)*; D: n = 18 WT, 18 *ser2prom3*::*deg-3(u662)*. Perfect turn is defined as a turn completed successfully without the male tail losing contact with the hermaphrodite. *t* test, black bars show mean, error bars show ± SD. *P* < 0.05 (*), *P* < 0.01 (**), *P* < 0.0001 (****).

Having observed that WT PVD enters into the tail rays, we sought to find whether this correlates with proficient mating ability. We found PVD branching into the tail rays precedes this behavior: L4 males, where PVD has not yet entered into the tail rays (Fig. 3 *L*), also lack a functional fan structure (16). This meant the earliest age we could analyze was young adults that have just molted, where PVD is already detected in some rays (Fig. 3 *L*). At this stage, many of the males displayed poor contact recognition (Fig. S10 *A*, YA, Movie S5, 9/45 showing no response at all). 24 h later, the same animals (kept in isolation) showed an improvement in mating turning and vulva recognition, comparable with same-aged virgin males (Fig. S10 *B* and *C*, Movie S5). Age therefore correlates with increased PVD entry into the tail rays (Fig. S10 *D*) and simultaneously contributes to mating performance (Fig. S10 *E*), although precise turning does not require a high percentage of rays to contain PVD (Fig. S10 *F*) nor directly correlates with a specific ray (Fig. S10 *G*). Following this observation, we used one-day adult males which show robust mating behavior in the WT, in order to compare backgrounds with altered PVD morphology.

### PVD affects male mating behavior

*mec-3(e1338), sax-7(dz156), mnr-1(dz175)* and *dma-1(tm5159)* show normal external tail morphology but no PVD ray extension (Fig. 3 *E-H*). This led us to wonder whether male behavior would be hindered in those circumstances. Indeed, all four genetic backgrounds impair male mating (Fig. S11), particularly in turning (Fig. S11 *E-H*). Combined, these results pointed to a potential role for PVD in a male-specific behavior, affected at least in part by its dendritic morphology.

As PVD may not be the only neuron affected in these backgrounds (32, 66, 69), we turned to a strain where PVD is specifically genetically ablated (25) by expression of *deg-3(u662)* dominant-negative mutation which induces neuron degeneration (in neurons which express DEG-3 nicotinic acetylcholine receptor subunit (70)) (Fig. 3 *K*). In the hermaphrodite, PVD genetic ablation results in altered crawling motion (25). We found that in males with PVD genetic ablation, mating behavior was impaired in all three steps of hermaphrodite contact recognition, turning and vulva stopping (Fig. 5). Together with the behavior impairment seen in the above four mutant backgrounds, these results suggest the PVD plays a role in adult male mating behavior, in particular during turning steps which require proprioception.

## Discussion

To our knowledge, although a neuron’s surrounding cells and connections may change in a sex-specific manner (41, 44, 71), very few examples in *C. elegans* exist where sex-shared neurons assume different dendritic morphology. A male-specific ectopic sprouting of the motor neuron PDB is one such example (46), with the sensory neuron PVD, as shown here, being the other. Our analysis revealed that while adult hermaphrodite PVD largely maintains its late L4 morphology (29, 30), male PVD undergoes further extension in the posterior region into the male-specific tail rays upon early adulthood (Figs. 1 *E* and *F* and *3 L*). This represents a morphogenesis mechanism distinct from that of a ‘resident’ ray neuron, RnB (Fig. 2). The ciliated RnB (56) is anchored at the ray tip (16) and assumes its shape by retrograde extension (23, 72) during the L4 stage ((16, 51), Fig. 2 *A*). In contrast, PVD does not express most ciliated neuron markers (58, 59) (*SI Appendix* Table S4, Figs. 2 *G* and *H* and S6), and extends into the rays only upon adulthood in an anterograde fashion (Fig. 3 *L*), producing intra-ray branching events which can be directly followed using time lapse imaging (Movie S6).

Our analysis supports a model where the male-specific PVD morphology relies on the sex-shared Menorin patterning complex (23), through ray SAX-7 expression (Fig. 4). In addition to its expression throughout the hypodermis (Fig. 4 *A* and *C*), SAX-7 was recently shown to be expressed in RnA neurons as well, affecting RnB morphogenesis (73). The defects observed in *sax-7(dz156)* mutant PVD, however, likely stem from truncation of the primary PVD branch prior to reaching the tail fan region (Fig. 3 *F*).

Based on electron microscopy analysis, we show PVD extends outside the primary neuron channel of the ray, and is embedded between the hypodermis and the structural cell (Figs. 3 *A* and *B* and *S9*). This placement is consistent with its reliance on hypodermal SAX-7 for correct morphogenesis (Fig. 4 *F-H*) and its ability to branch and U-turn within each ray (Figs. S7 and S8, Movie S8), and additionally implies it is not open to the external environment. While our results suggest PVD and RnB are separate in the tail fan region (Fig. 2), we note EM-based connectome studies detected synapses between PVD and RnB in the pre-anal ganglion region (3).

In addition to developing a male-specific dendritic morphology, it is possible that sexually-dimorphic connectivity also occurs during early adulthood, in parallel with an improvement in mating behavior (Fig. S10 *A-C*). While PVD-PVC connectivity was followed in the hermaphrodite (74), it is important to note that much of our knowledge of PVD synaptic partners relies on a male EM series reconstruction (N2Y), for which comparable hermaphrodite reconstructions are as yet missing (2, 3).

Alongside ‘dedicated’ ray neurons such as RnA (68), we show that PVD is important for male contact response, turning around the hermaphrodite and vulva recognition (Fig. 5). Importantly, *sax-7* mutant males were previously shown to have defects in contact response as well as vulva location efficiency (66). While in our hands a turning defect was the most pronounced, this may stem from the different *sax-7* allele utilized as well as different behavioral scoring criteria (see Materials and Methods). Recently, mutants of *kpc-1* (encoding a Furin enzyme) were also shown to have male mating defects alongside impaired PVD arborization in the tail region (75). Our results provide evidence for additional male-specific behavioral defects (Fig. S11) in animals with impaired PVD morphology (Fig. 3 *E-H*). While we cannot rule out the possibility that PVD branching in certain ray combinations is important for turning behavior, as was demonstrated by global ablations of all or some ray neurons (48, 68), this is difficult to ascertain in WT adults (Fig. S10 *G*). PVD branching into the tail rays precedes a striking change in male mate-seeking behavior and mating capacity from L4 to adulthood (Fig. S10 *A-D*). While PVD activity and morphology are not the sole contributor of male sexual behavior (48, 68, 76), PVD may be modulated as part of a male-specific, adult-specific, learning process. Complete and specific ablation of the PVD results in male behavior defects (Fig. 5), which are consistent with PVD’s role as a proprioceptor (25, 26), and extend the role of PVD in males.

Our results uncover a previously uncharacterized role for PVD in male mating behavior, and reveal a sexually dimorphic instance of post-larval specific branching into the male tail rays, which seems integral to the early adult dendritic arbor. Further work is required to directly establish the mechanisms regulating PVD ray entry, and to elucidate the direct function of this dendritic morphology together with body candelabra in proprioception and male behavior.

## Materials and Methods

### Strains and their maintenance

A complete list of strains used in this study is found in the *SI Appendix* Table S1.

Worms were kept at 20°C unless specified, on NGM plates with 150 µl OP50 *Escherichia coli* grown overnight in LB medium, using standard methods (78). Males which are not from a *him-5(e1490)* genetic background were obtained by subjecting L4 hermaphrodites to 32°C for obtaining a higher incidence of spontaneous males (15), which were then propagated with non-heat-treated hermaphrodites. Additional details related to synchronization, fluorescent marker expression profile, and *eff-1* growth rates are found in *SI Appendix*.

The *ser2prom3::GFP* marker labels the sensory PDE neuron as well, whose cell body is adjacent to PVD (31, 77) PDE sends an anterior but also a posterior ventral process (2), observable in males, but also hermaphrodites, as elegantly shown by O’Brien et al., (31). It can be seen most clearly in Fig. 3 *E* (green arrow), but can also be seen in Fig. 3 *D* and Fig 1 *D* and *E*.

### Transgenic lines

Transgenic animals were obtained by microinjection (79). PVD feminization was analyzed in two separate transgenic lines. Hypodermal *dpy-7p::sax-7(s)* rescue experiments analyzed two different transgenic lines as one-day adults (L4+1 day).

### Electron microscopy

Electron microscopy was performed as described in (49). Details of the sections used are found in

*SI Appendix*.

### Mating assays

Mating was scored following Liu and Sternberg, (48), and experiments set up as published previously by Oren-Suissa *et al.,* (41). In our scoring of vulva recognition, we score any stopping or hesitation of the tail in the vulva region as a successful vulva recognition event. This is in contrast to the precise methodology used by Kim and Emmons, 2017 (66), which takes into account only complete stops with a duration of 10 seconds. Further details are found in *SI Appendix*.

### Imaging

Worms were typically immobilized in 0.05% tetramisole (Sigma T1512) in M9 buffer (78). Briefly, a 3% agar pad was prepared (Bacto-Agar in DDW) and worms were loaded onto a 4 µl drop of tetramisole placed directly on the pad. A coverslip was then secured with thin strips of marking tape. Some worms were similarly mounted on 3-5% agarose using 30 mM sodium azide (Sigma S2002) in M9; additional details related to animal immobilization and the microscopy system utilized are found in *SI Appendix*.

### Neuron tracing

Neuron tracing was performed on three-dimensional stacks semi-manually by the Fiji plugin SNT 3.2.3 (Plugins → Neuroanatomy → SNT) (80) and colored manually within the software. Rotation videos of such tracings were obtained using the software’s 3D viewer. Cell placements (*e.g.* RnB cell bodies) were traced manually using the Fiji Cell Counter plugin on a three-dimensional image stack, subsequently projected to the XY plane. For figure clarity, this projection was then manually traced over in a thicker stroke. Additional details are found in *SI Appendix*.

### PVD-specific genetic ablation and interacting neurons

Additional details related to the specificity validation of PVD genetic ablation by *ser2prom3::deg-3(u662)* in males are found in *SI Appendix*

### Image analysis and quantification

Analysis of male tail rays for neuron presence was manually evaluated using Fiji (ImageJ, NIH) by merging DIC and fluorescent channels and following the fluorescent signal through the Z-series in three dimensions. Rays which were not clearly visible by DIC were not scored. Most animals were captured in a lateral view; the distal PVD neuron was typically difficult to analyze, hence some animals have data for one side but not the other. Colocalization of PVD and RnB was followed by scanning through the Z-series of the two merged channels, and length measurements were manually taken using a Segmented Line tool in Fiji. The figures presented are Z-axis projections, typically with DIC in gray and fluorescent signal pseudo-colored cyan. Transgenic lines were imaged alongside non-transgenic siblings and analyzed blindly.

Images were edited for brightness, contrast, correct scaling and channel merging using Fiji (imageJ, NIH). Images of neurons are maximal-intensity projections across a Z-series stack, while DIC images are a single Z position. Figures were prepared using Adobe Illustrator CS5 (Adobe Inc.). Plots were prepared using Prism 9.0.2 (GraphPad).

Analysis of PVD orders was performed on image series taken around or just anterior to the cell body, by manual annotation using the Fiji Cell Counter plugin (Plugins → Analyze → Cell Counter → Cell Counter) on a three-dimensional image stack. Different counter types were renamed for each branching order, and final counts were manually copied. ‘Ectopic’ branches were defined as processes which deviate from the four-order candelabrum shape, *i.e.* endpoints which are not quaternary ‘candles’ (81).

### Statistics

All statistical tests were performed using Prism 9.0.2 (GraphPad). Unless specified otherwise, *p* values are represented in asterisks as follows: *P* <0.05 (*), *P* <0.01 (**), *P* <0.001 (***); *P* <0.0001 (****). Specific tests are described individually for each figure.

## Figure preparation

Final figures were prepared using Illustrator CS5 (Adobe Inc.). Movies were compiled, annotated and saved using Fiji (ImageJ, NIH).

## Acknowledgments

We are grateful to Yehuda Salzberg, Hagar Setty and Meital Oren for strains (EB1564, EB1271, EB1653, DA509, MOS81, MT5383, PT2565), DNA constructs (pMO32) and valuable advice regarding mating experiments, to Kang Shen for strains (TV15911, TV15916, TV19321) and DNA constructs (pXD22), to Millet Treinin for MF288 and to Arantza Barrios for PS3380 and EM1106. We thank Scott Emmons, Hannes Bülow, Oliver Hobert and Maxwell G. Heiman for their valuable comments regarding PVD connectivity and ciliated neuron hallmarks. Son Le Tho for preliminary analyses of sexual dimorphism in PVD branching. Shay Stern, Dan Cassel, Meital Oren and the

Chalfie Lab members for critically reading the manuscript. Some strains were obtained from the *Caenorhabditis* Genetics Center (CGC) which is funded by the NIH Office of Research Infrastructure Programs (P40 OD01440). Research in our lab has received funding from the Israel Science Foundation (2462/18, 2327/19, 178/20 and 1575/21 to B.P.) and Dirección General de Asuntos del Personal Academico, Programa de Estancias de Investigación (PREI), UNAM, Mexico City (B.P.). Electron microscopy was supported by NIH OD R24 010943 (D.H.H.).

