## Supplemental Information for "The PVD neuron has male-specific structure and mating function in *C. elegans*"

**This PDF file includes:**

Supporting text
Figures S1 to S11
Tables S1 to S4
Legends for Movies S1 to S6
SI References

**Other supporting materials for this manuscript include the following:**

Movies S1 to S6

### Supporting Information Text

#### Strains and DNA constructs

**Strains.** The complete list of strains used in this study is found in **Table S1**.

**DNA constructs**. The complete list of DNA constructs used in this study is found in **Table S2**.

#### Additional details for Materials and Methods

**Electron microscopy.** The transverse section used in this work belongs to the same animal and same ray depicted in Fig. 2C of (1), which shows a more distal section, in which PVD is no longer visible. The PVD was followed over several sections and is distinct from the ray neurons RnA and RnB, tapering towards the end of the ray and generally thinner than RnA and RnB. It has a clear cytoplasm, elongated outline when viewed from the side, or closer to circular when viewed in cross-section, and few to no microtubules. From the sections observed, PVD tends to run inserted between the hypodermis and the structural cell.

**Mating assays.** When scoring turning behavior, a 'perfect' turn was defined as a turn completed successfully without the male tail losing contact with the hermaphrodite (see 'good turn' vs 'sloppy turn' in Fig. 4 (2)). Contact response was annotated as a reversal with tail pressing against the hermaphrodite immediately following hermaphrodite tail contact. Vulva recognition was annotated as any stopping around the vulva region, and not constrained by time. L4-stage males (wild type and test group) and *unc-31(e928)* hermaphrodites were picked a day prior to the experiment, kept separately. NGM plates were seeded with a single drop (~10 µl) OP50. The following day (approximately 16 h later), with the OP50 dried to a matte finish, 15 hermaphrodites were transferred onto a plate. Male plates (wild type and test group) were coded and then blinded by a second person, which had no knowledge of their genotype. For each experiment, a single male animal was placed just outside the OP50 spot using an eyelash attached to a toothpick on the plate and monitored under a dissecting scope for 15 minutes from the first tail contact of a hermaphrodite, or until mated (if sooner). Animals which did not approach within one body-length away from a hermaphrodite after five minutes were repositioned with an eyelash. Mated hermaphrodites were replaced. For some experiments, animals which displayed difficulty with spicule insertion, a step which may depend on the hermaphrodite used (3), were assayed on a second hermaphrodite for an additional 15 minutes or until mated, typically by removing the first hermaphrodite and allowing the male to locate a second mate on the plate. Cases where two hermaphrodites were used were pooled together to collect the overall male behavior in both attempts. Males were assayed alternating between both blinded test groups to avoid a batch effect due to different age or plate condition. Assays for young adult-to-adult transition utilized a single hermaphrodite per male. All experiments involved manual annotation of the mating sequence *in situ*. Most experiments were recorded using a Hamamatsu Orca-ER EMCCD camera, mounted on a Zeiss SteREO Discovery.V8 dissecting scope, using Micromanager 2.0 (ImageJ, NIH), set a 2 x 2 binning, imaging 1000 frames per 04:15 minute video; several such videos were captured for each mating. Spicule insertion and sperm transfer were definitively established for most of the experiments. Mating efficiency (number of progeny sired (4)) was not determined.

Importantly and as noted in (5), animal mounting (see 'Imaging') utilized tetramisole (levamisole), which induces spicule protraction. This effect was at times irreversible and led to mating defects in cases where the animals were re-evaluated (data not shown). To mitigate this, these animals were closely observed prior to mating contact, and were censored if the spicules were protracted. The improved behavior of young animals, assayed twice (which did not show spicule extension) indicates that the mounting, imaging and recovery process by itself does not significantly harm male mating behavior.

**Imaging and animal mounting.** Worms were immobilized in 0.05% tetramisole (Sigma T1512) in M9 bufferTetramisole (6, 7). In some cases and for brief imaging sessions where animals were not recovered, animals were similarly mounted in 30 mM sodium azide (Sigma S2002) in M9 on 3% agar or 5% agarose (Lonza #50004). This was used to minimize animal movement when imaging strains with both RnB and PVD markers (*bxIs14; dzIs53*) and also utilized with some 2-day adult BP2285 *wyIs592; him-5(e1490)* and with BP2297 *ddIs290; him-5(e1490)*. No difference was noted in PVD presence in tail rays among same-aged two-day adult WT animals mounted in tetramisole or sodium azide (n = 10,11 respectively): 78.8±14.1 vs 81.3 ± 13.5, Mean± Std, respectively. Male PVD morphology in L4 WT is not observably altered by the use of tetramisole compared with sodium azide mounting, although thin quaternary branches are somewhat clearer in azide mounting (data not shown).

Animals were imaged using one of two microscope systems: 1) an inverted confocal microscope (Nikon Ti Eclipse) with a CSU-X spinning disk unit (Yokogawa), and either a Plan-Fluor oil 40X NA 1.3 or an Apo oil 60x NA 1.4 (Nikon) objective with or without an added 1.5x magnification, capturing serial z-axis sections spaced 0.6 µm apart captured by an iXon3 or iXon Ultra EMCCD camera (Andor, Belfast), using Metamorph (version 7.8.1.0, Molecular Devices); 2) an inverted confocal microscope (Nikon Eclipse Ti-2) with a CSU-W1 spinning disk unit (Yokogawa) equipped with a water Plan-Apochromat 60X NA 1.2 objective (CFI) capturing serial z-axis sections spaced 0.3 µm apart using a Prime BSI sCMOS camera (Teledyne Photometrics) through NIS-Elements AR interface software (Nikon). GFP and Kaede were imaged using 488 nm laser excitation, mCherry and photoconverted Kaede were imaged using 561 nm laser excitation. Kaede photoconversion was performed by 405 nm laser excitation. Experiments where both PVD neurons were imaged involved opening the slide, adding a drop of M9 buffer, repositioning the animal using an eyelash attached to a toothpick, and replacing the coverslip. Animals which were to be imaged again at a later time point were similarly removed from the slide and placed to recover on an NGM plate with OP50 as described.

**PVD-specific genetic ablation and interacting neurons.** While the RnB-specific *pkd-2::GFP* used does not mark R6B (8–10), a cross of the genetic ablation strain into a *pkd-2::GFP* strain reveals all other RnB neurons are unaffected. While we cannot directly establish whether R6B is affected by the *ser2prom3*::*deg-3(u662)* construct through gap junction interactions, no vacuolated neurons were observed by DIC in the lumbar ganglion. Additionally, while killing of electrically-coupled neurons to those expressing *deg-3(u662)* has been hinted (11), the PVD is strongly associated with the CAN neuron (12, 13), which, although its precise function remains unclear, has a lethal rod-shape phenotype when ablated (13, 14), which was not observed.

No data was found regarding *deg-3* expression in male ray neurons, however hermaphrodite RNA sequencing data is available at the single neuron level (https://cengen.shinyapps.io/CengenApp/ ; (15)). This data supports the hypothesis in (16), which the OLL neuron pair, the only other observable neuron expressing a *ser-2prom3* reporter (16, 17), is not degenerated in *ser2prom3*::*deg-3(u662)* animals since it expresses ~100-fold less *deg-3* (https://cengen.shinyapps.io/CengenApp/; (15)). Based on hermaphrodite data, body muscles express similarly low levels of *deg-3* (https://cengen.shinyapps.io/CengenApp/; (15)). This tentatively supports the notion that muscles are likely unaffected by ectopic *deg-3(u662)* even if low expression of *ser2prom3* is present (a different *ser-2* promoter region, *ser2prom1*, is expressed in a sexually-dimorphic fashion in male posterior muscles, including the diagonal muscles used in mating (17)).

***eff-1* mutant animal growth rates.** Development timing assessment of WT animals was obtained from (18, 19), yielding roughly consistent ratios between 16°C, 20°C and 25°C, with roughly 25% of pre-adult development spent as L4 (26±1.7%) (19). Development time at 16°C can be corrected to an equivalent 20°C dividing by 1.6 and 25°C corrected to 20°C dividing by 0.76 (19). We used 16°C values for growth at 15°C (correction factor 1.6), and the mean of 20°C and 25°C for growth at room temperature (22-23°C) (correction factor 0.87). *eff-1* animals grow somewhat slower than WT (20), and the null mutant *ok1021* at 20°C seemed to reach young adulthood after 72 h. Based on this time point and assuming the proportion of time spent at each developmental stage is WT-like, we obtained an approximate development rate in hours, found in **Table S3**. These values were used to convert real hours of development to their (hypothetical) 20°C equivalent by the correction factors mentioned above.

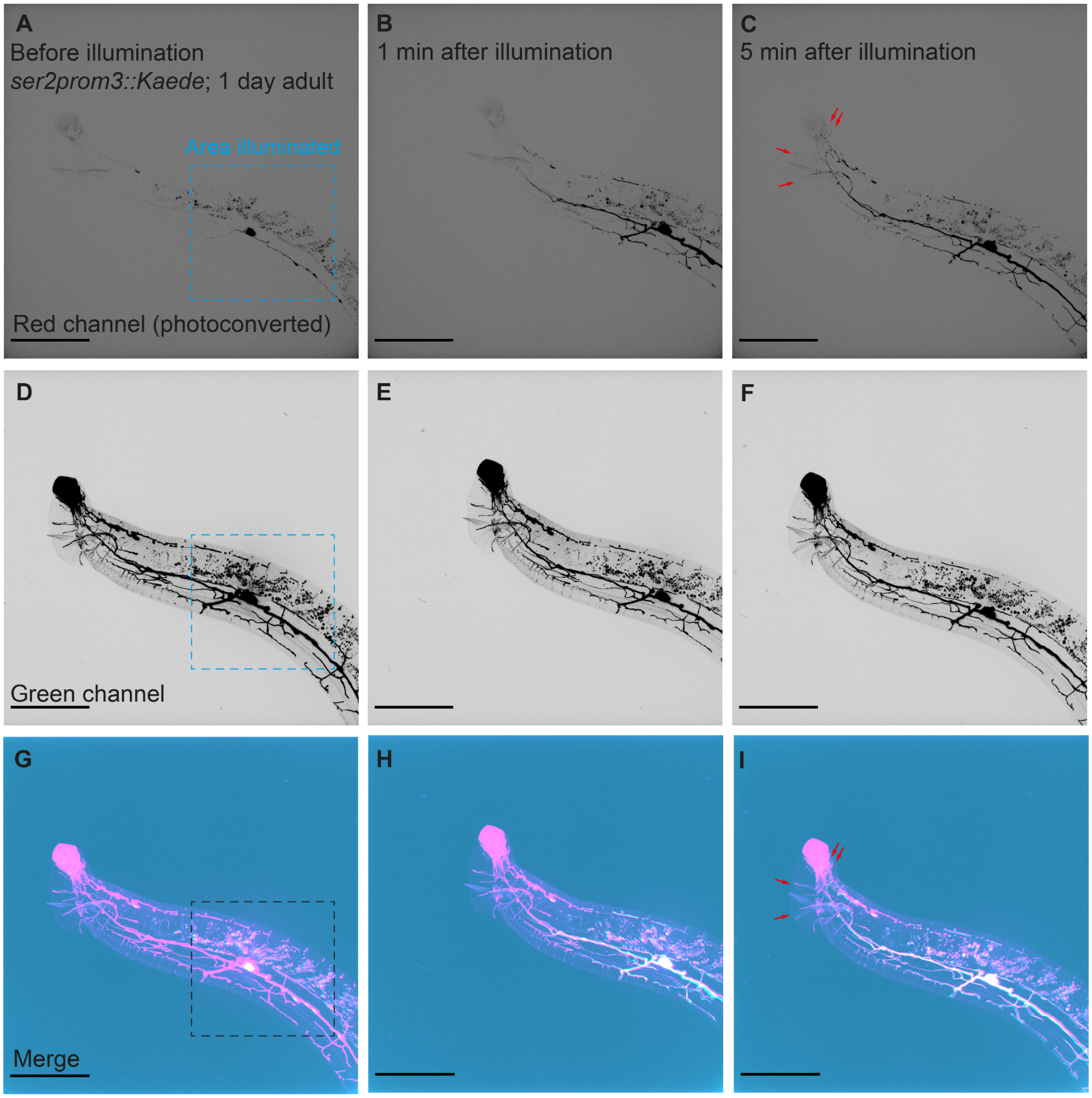

**Fig. S1. Kaede photoconversion supports PVD entry into tail rays**
(A) Red channel *ser2prom3::Kaede* PVD signal prior to photoconversion (negative image); dashed blue frame indicates approximate area illuminated to achieve Kaede photoconversion from green channel to red channel emission.
(B) Red channel *ser2prom3::Kaede* PVD (negative image) 1 min after photoconversion.
(C) Red channel *ser2prom3::Kaede* PVD (negative image) 5 min after photoconversion; red arrows, signal in rays.
(D-F) Respective green channel *ser2prom3::Kaede* PVD signal taken with panels A,B,C, respectively.
(G-I) Merged channels A,D; E,H; F,I, respectively. For clearer contrast, the green channel is pseudocolored magenta, while the weaker red channel is pseudocolored cyan; white signal indicates overlap between both channels. Scale bar is 50 μm.

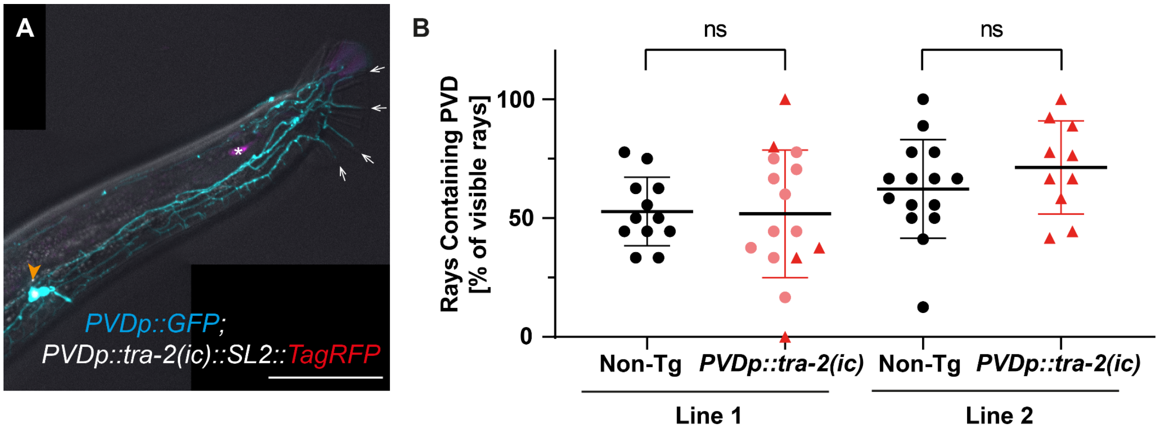

**Fig. S2. PVD tail ray morphology is independent of PVD-specific feminization**
(A) One-day adult male expressing *tra-2* constitutively active intracellular domain (*ser2prom3::tra-2(ic)::SL2::2xNLS-TagRFP*) inducing a hermaphrodite fate (21). PVD marked by *ser2prom3::GFP*. Arrows point to PVD branching in the rays, orange arrowhead marks PVD cell-specific RFP expression. Asterisk marks *podr-1::RFP* expression unrelated to the transgene.
(B) Quantification of *ser2prom3::GFP* PVD entry into the rays of the male tail in two independent transgenic lines [*ser2prom3::tra-2(ic)::SL2::2xNLS-TagRFP 20 ng/µl pmyo-2::GFP 10 ng/µl pBluescript empty vector 70 ng/µl*] in transgenic versus non transgenic (non-Tg) sibling animals (determined by *pmyo-2::GFP*). Red triangles denote animals with an additional clear PVD RFP signal, not observed in any of the controls. Unpaired *t* test, ns = not significant ($p>0.05$). Black bars show mean, error bars show ± SD.

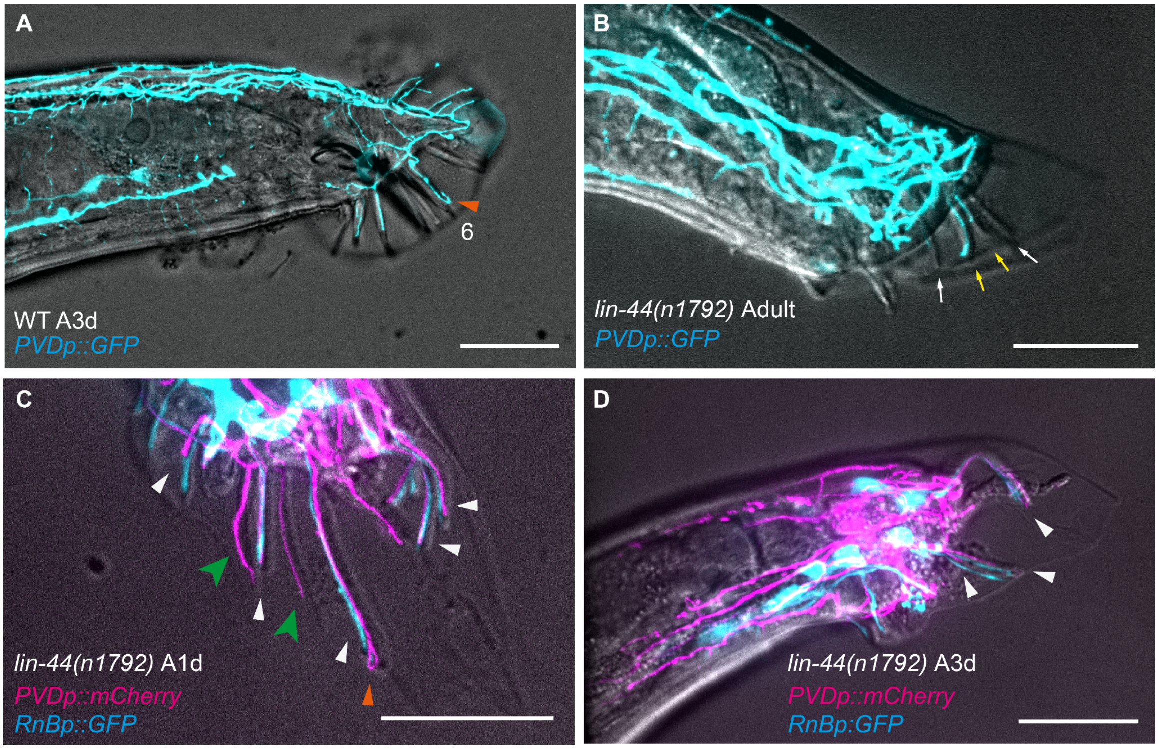

**Fig. S3. PVD processes can enter *lin-44* (Wnt) mutant tail rays, independently of RnB presence**
(A) Dorso-lateral view of WT *him-5(e1490)* expressing *ser2prom3::GFP* PVD marker with an example of PVD U-turn within ray 6 (orange arrowhead). Scale bar is 25 μm.
(B) *lin-44(n1792); him-5(e1490)* adult expressing *ser2prom3::GFP* PVD marker. Yellow arrows indicate a ray with PVD signal, white arrows indicate rays without PVD signal. Scale bar is 25 μm.
(C) *lin-44(n1792); him-5(e1490)* 1-day adult (A1d) expressing *pF49H12.4::mCherry* PVD marker and *pkd-2::GFP* RnB marker. White arrowheads point to non-overlapping PVD and RnB intra-ray signals, green arrowheads point to PVD entering a ray in the absence of RnB (one of which is possibly ray-6-like), orange arrowhead marks U-turn of a PVD branch. Scale bar is 25 μm. Of n = 11 sides observed, one showed two rays lacking *pkd-2::GFP*. Three additional examples could be a WT-like absence similar to R6B.
(D) *lin-44(n1792); him-5(e1490)* 3-day adult (A3d) coexpressing *pF49H12.4::mCherry* PVD and *pkd-2::GFP* RnB markers. White arrowheads point to non-overlapping PVD and RnB intra-ray signals. Scale bar is 25 μm.

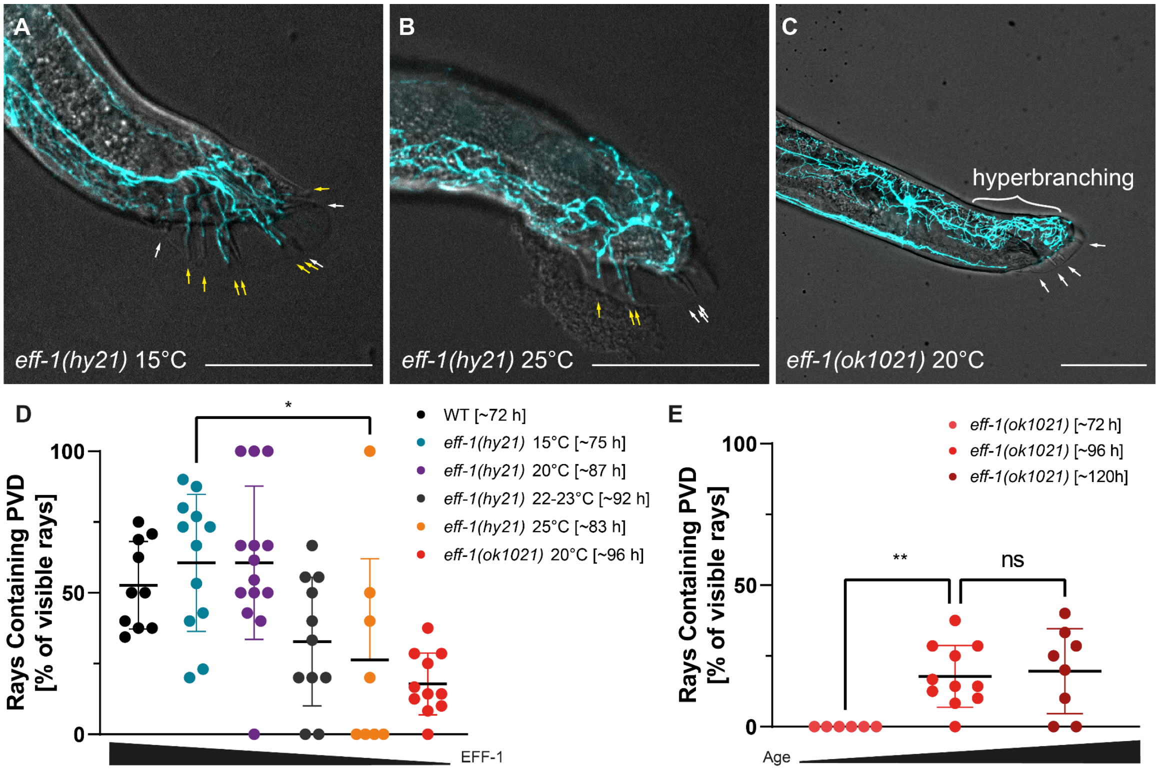

**Fig. S4. EFF-1 dosage affects PVD ray entry**

(A) Merged DIC (gray) and PVD (cyan) of an L4 + 24 h temperature-sensitive *eff-1(hy21); him-5(e1490); ser2prom3::GFP* animal raised at the permissive temperature (15°C). Yellow arrows indicate rays with PVD signal, white arrows indicate rays without PVD signal. Scale bar is 50 µm.
(B) Merged DIC (gray) and PVD (cyan) of an L4 + 24 h temperature-sensitive *eff-1(hy21); him-5(e1490); ser2prom3::GFP* animal raised at the restrictive temperature (25°C). Yellow arrows indicate rays with PVD signal, white arrows indicate rays without PVD signal. Scale bar is 50 µm.
(C) Merged DIC (gray) and PVD (cyan) of an adult (four days from embryo, approximately L4 + 24 h) null mutant allele *eff-1(ok1021); him-5(e1490); ser2prom3::GFP* animal raised at 20°C; note hyperbranched phenotype in the dorsal side (brackets); white arrows indicate rays without PVD signal. Scale bar is 50 µm.
(D) Quantification of PVD dendrite extension into rays in *eff-1* mutant animals (background *him-5(e1490); ser2prom3::*GFP). Timing is corrected to equivalent hours at 20°C (see Materials and Methods Table S1, n = 10, 12, 14, 11, 8, 11, respectively. One-way ANOVA comparing 72 h to 96 h and 96 h to 120 h; Šídák multiple comparison correction. Black bars show mean, error bars show ± SD, except 25°C, showing + SD. $P < 0.05$ (*), $P < 0.01$ (**), $P < 0.001$ (***); $P < 0.0001$ (****). WT representative data is identical to WT A1d in figure 2I.
(E) Quantification of PVD entry into the tail rays in *eff-1(ok1021)* null mutants of different ages (background *him-5(e1490); ser2prom3::GFP*). ~96 h data is replicated from panel **I**; n = 6, 11, 8 respectively. One-way ANOVA; ns ($P > 0.033$) unless specified; Šídák multiple comparison correction. Black bars show mean, error bars show ±SD. $P < 0.05$ (*), $P < 0.01$ (**), $P < 0.001$ (***); $P < 0.0001$ (****).

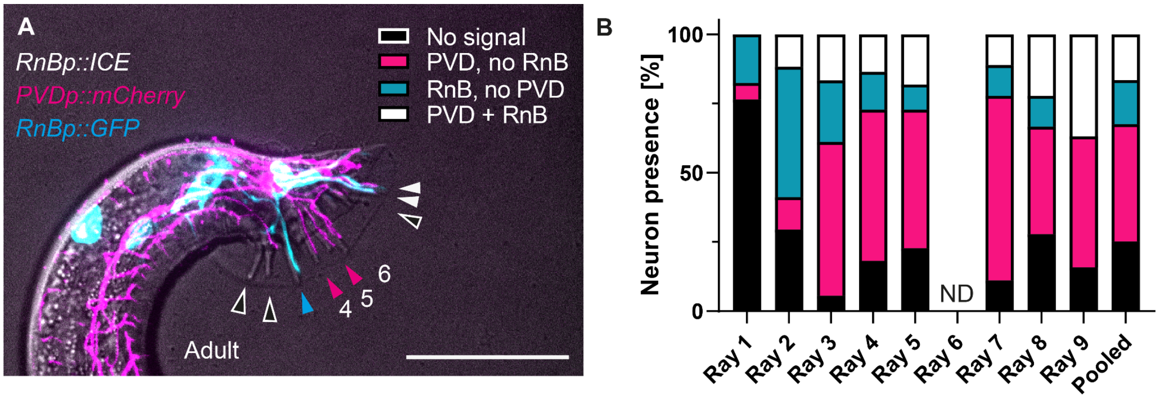

**Fig. S5. RnB presence is not required for PVD entry into the tail rays**
(A) Adult *him-5(e1467)* expressing *pkd-2::ICE* human caspase in RnB as well as *pkd-2::GFP* RnB marker and *pF49H12.4::mCherry* PVD marker. Black arrowheads mark rays without RnB or PVD signal, magenta marks PVD presence in the absence of RnB, cyan marks RnB without PVD, white arrowheads point to non-overlapping PVD and RnB intra-ray signals. Rays 4, 5, 6 are indicated for orientation. Scale bar is 50 μm.
(B) Percentage of rays observed showing the presence of none, PVD, RnB or both, in adult *him-5(e1467)* expressing *pkd-2::ICE* human caspase in RnB, in males expressing *pkd-2::GFP* RnB and *pF49H12.4::mCherry* PVD markers. Color annotation as in panel (A). Ray 6 is not scored since *pkd-2::GFP* is absent regardless of ICE expression (8). n = 17, 17, 18, 22, 22, 18, 18, 19 and 151 pooled rays examined, respectively. Fisher's Exact $p = 0.22$ for pooled counts (No PVD, no RnB: 38; PVD, no RnB: 64; No PVD, RnB: 24; Both: 25).

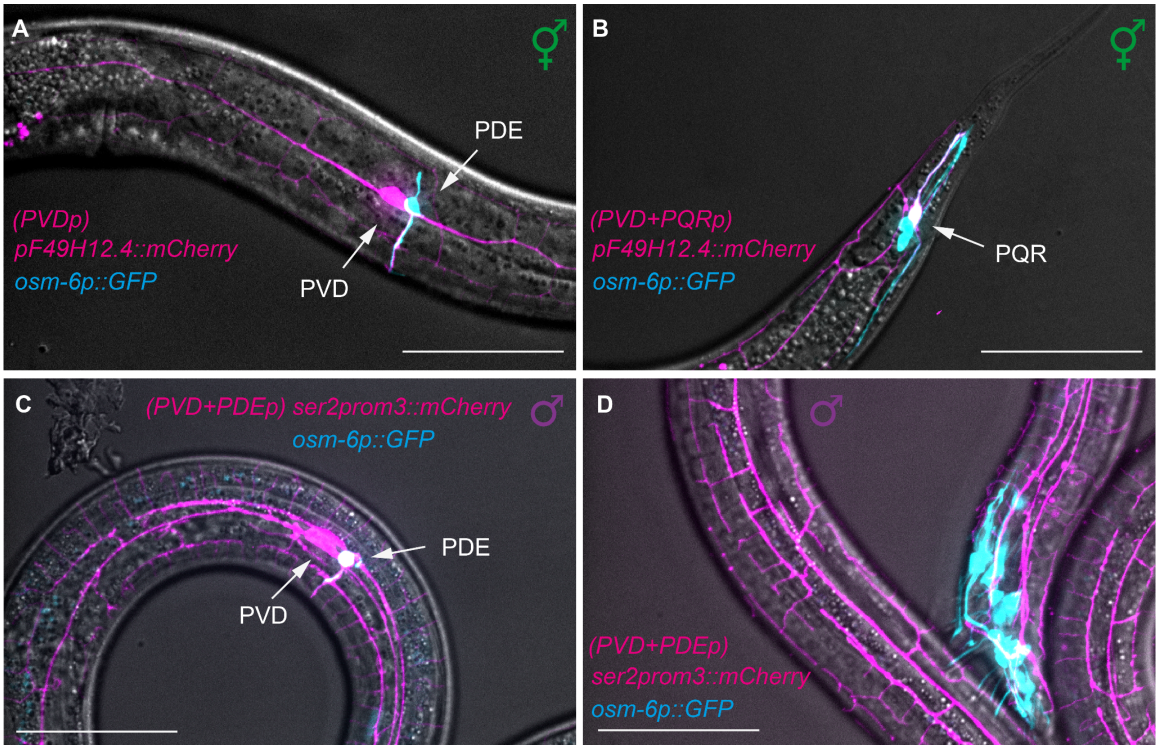

**Fig. S6. *osm-6p::GFP* ciliated neuron marker is not expressed in PVD**
(A) PVD cell body region of an adult hermaphrodite expressing *osm-6p::GFP* (cyan), expressed in ciliated neurons including PDE and PQR (22, 23), and *pF49H12.4::mCherry*, expressed in PVD (magenta). Scale bar is 50 μm.
(B) Tail region of an adult hermaphrodite expressing *osm-6p::GFP* (cyan), expressed in ciliated neurons including PDE and PQR (white cell body and process), and *pF49H12.4::mCherry* (magenta), expressed in PVD and PQR. Channel signal colocalization appears white. Scale bar is 50 μm.
(C) PVD cell body region of a young adult male expressing *osm-6p::GFP* (cyan), expressed in ciliated neurons including PDE and PQR, and *ser2prom3::mCherry* (magenta), expressed in PVD and OLL (24). 15/15 adult animals show colocalization of the two markers in PDE, but not PVD, cell bodies. Channel signal colocalization appears white. Scale bar is 50 μm.
(D) Tail region of a young adult male expressing *osm-6p::GFP*, expressed in ciliated neurons including RnAs, RnBs, PHA and PHB (cyan) , and *ser2prom3::mCherry*, expressed in PVD (magenta). Scale bar is 50 μm.

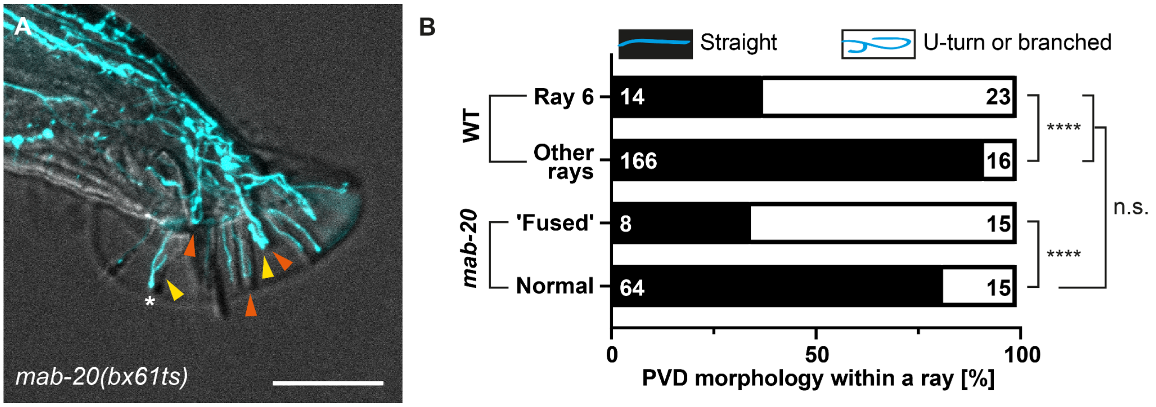

**Fig. S7. Ray width affects PVD propensity for U-turns and intra-ray branching**
(A) *mab-20(bx61ts); him-5(e1490)* adults expressing *ser2prom3::GFP* PVD marker, raised at 20°C. Orange arrowheads indicate PVD U-turns; yellow arrowheads mark branched PVD within a ray; asterisk marks 'fused' ray. Scale bar is 25 μm.
(B) Effect of ray width on PVD tendency to branch or turn within a ray (white) compared to straight PVD branch (black); comparing WT (*him-5(e1490); ser2prom3::GFP* PVD marker) ray 6 with all other rays in a 2-day adult animal and 'fused' rays with normal rays in *mab-20(bx61ts); him-5(e1490); ser2prom3::GFP* PVD marker 1.5-day adult animals. The number of rays is shown within the bars; X axis presents this data as percentage. Statistics were performed using Fisher's exact test using the number of rays. Comparing all WT rays with all non-'fused' *mab-20(bx61ts)* rays is not statistically significant. $P$ <0.0001 (****).

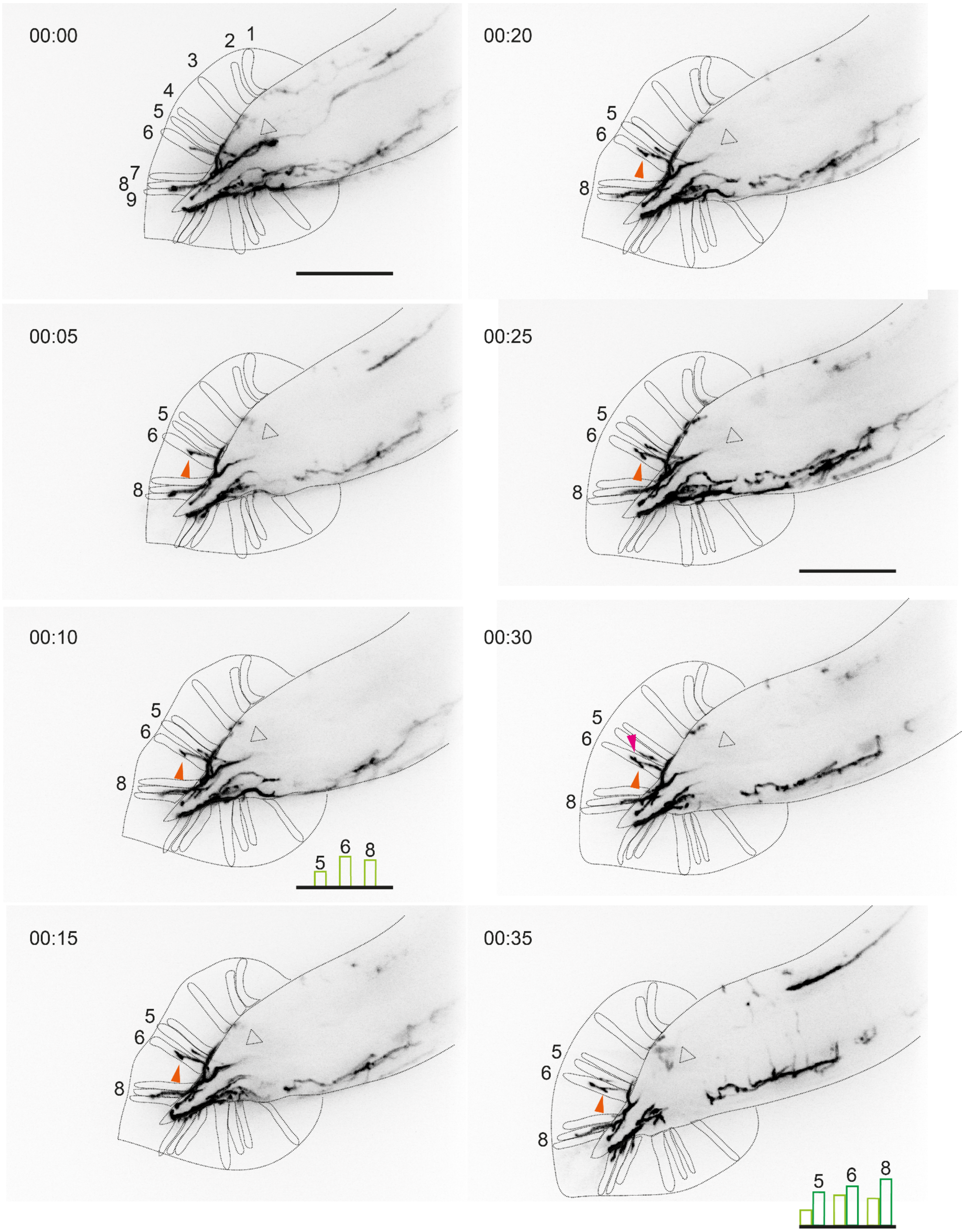

**Fig. S8. Timelapse of WT young adult PVD growing within the rays**
Negative images from a timelapse in minutes highlighting two growth events of ectopic processes in ray 6 (orange and magenta arrowheads). Green bars demonstrate approximate growth in the PVD processes of rays 5, 6 and 8 between 00:10 (light green) and 00:35 (dark green), as measured based on the projected image. Animal is a young adult WT (a few hours after molting) *him-5(e1490)* male expressing *ser2prom3::GFP* PVD marker mounted in 0.05% tetramisole on a 5% agarose pad. Signal is inverted. Scale bar is 25 μm

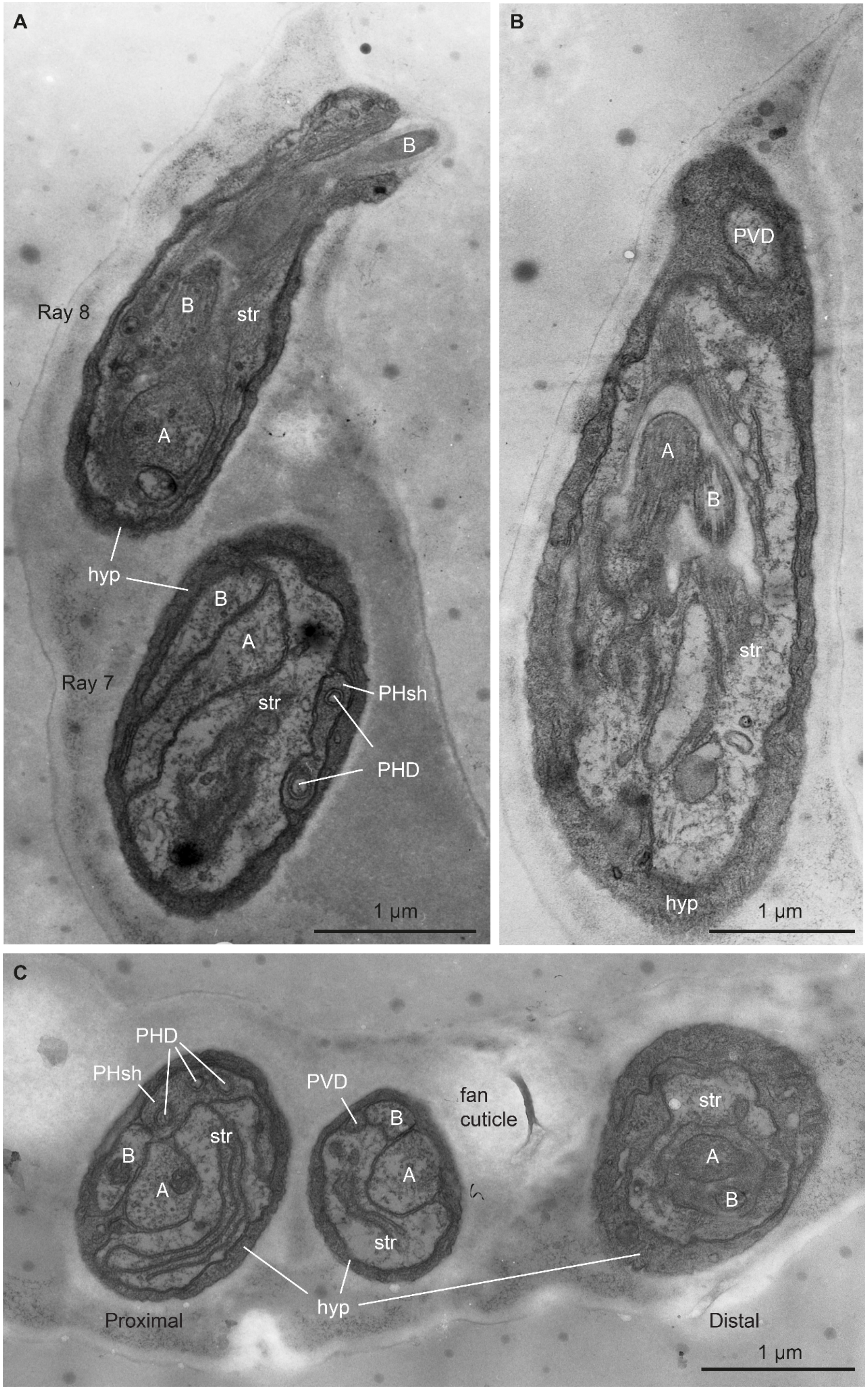
A
B
Ray 8
B
str
A
hyp
B
A
Ray 7
str
PHsh
PHD
1 µm
B
PVD
A
B
str
hyp
1 µm
C
PHD
PHsh
PVD
fan
cuticle
B
str
B
A
A
str
str
A
B
hyp
Proximal
Distal
1 µm

**Fig. S9. EM sections show PVD is outside the ray neuron channel**

(A) The tip of ray 8 contains only two ciliated neurons. Distal view of ray 8 at the ray opening on the ventral side of the fan cuticle. R8B (B) is observed in two cross-sections within the same slice, one where the cilium tip exits the ray, and another view where the cilium sits inside the channel created by the structural cell (str). The cilium of R8A (A) lies next to R8B in the channel, and has a much larger diameter. The nearby view of ray 7 in cross-section is midway along the ray, prior to the dendrites of R7A and R7B entering the str channel (no channel has formed yet, but the str process has folded membranes that can expand to form the channel). Two thin fingers of the PHD cilium are seen here, wrapped in a process from the phasmid sheath cell (PHsh). PHD fingers can occasionally enter from the base of several rays (probably 6 through 8). The cilia of PHD are not known to enter a ray, only these fingers. PVD is not observed.

(B) Distal view of a ray in lengthwise view reveals a PVD process which reaches the distal edge of a ray, beyond the cilia of RnA (A) and RnB (B), which both lie inside the structural cell (str) channel. From this viewing angle, it is not possible to determine if the PVD process will form a U-turn. The entire ray is always surrounded by a continuous cylinder created by the local hypodermal cell (hyp), as well as by the fan cuticle.

(C) Cross-section of the male fan shows three adjacent rays lying side by side, wrapped by the fan cuticle. The more proximal ray on the left is closer to its base, and contains a prominent structural cell process (str), two ray neuron dendrites, RnA and RnB (A and B, respectively), and several thin fingers from the PHD cilium, which is wrapped in a process from the nearby phasmid sheath cell. There is no evidence for a PVD dendrite here, and the viewing angle excludes this as a possibility. The middle ray holds four processes (str, A, B, PVD) wrapped in a cylinder of hypodermal (hyp) tissue. The PVD process is markedly smaller than all other elements. PHD was not recognized as a separate neuron until recently (25), and has not been shown to extend inside a ray before these examples. The ray on the right is more distal, and here exactly two ray neuron cilia (RnA and RnB) have entered the expanded structural cell channel (str). PHsh: phasmid sheath cell.

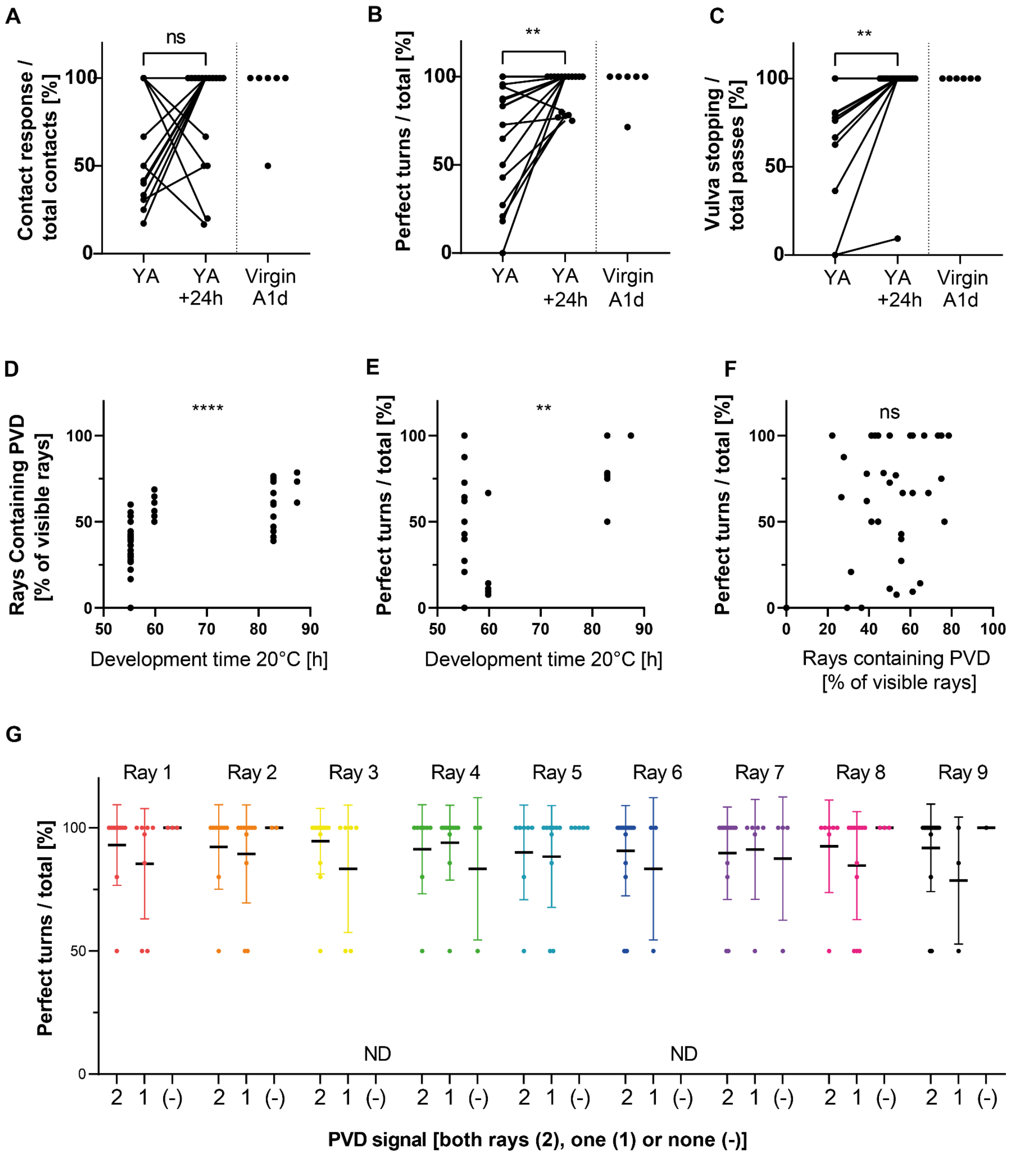

**Fig. S10. Ray PVD extension precedes male mating behavior onset.**

(A-C) Hermaphrodite recognition, turning and vulva recognition, respectively, in WT *him-5(e1490); ser2prom3::GFP* PVD marker virgin animals assayed shortly after molting to adulthood (young adults, YA) and upon re-evaluation 24 h later (YA+24h); lines connect the same individual. Six virgin one-day adult (A1dA) males were assayed in parallel to young adults (YA), to confirm previous mating experience is not required for the age-dependent improvement noted. Statistics was performed by a Wilcoxon matched-pairs signed rank test, and between YA+24h and unmated virgin 1dA by a nonparametric Mann-Whitney test. n =16 animals behaving on both timepoints. $P$ <0.05 (*), $P$ <0.01 (**).

(D-F) Individual plane views of PVD-age, behavior-age and behavior-PVD, respectively. G: n = 52, of which 35 YA and 17 assayed again as YA + 24h. H,I: 25 YA and 15 YA+ 24h. The two groups compared by *t* test are shown as circles and squares, as shown under each panel. Nonparametric Spearman correlation, ns: not significant, $P$ <0.005 (**); $P$ <0.0001 (****).

(G) The presence or absence of PVD branching within individual rays is not correlated with decreased turning ability. N =23 animals, four missing data for rays 8+9. No statistical analysis was performed since very few examples of PVD absence were found. Both rays in a pair: 2; one of a pair: 1; none: (-). Black bars show mean, error bars show ±SD.

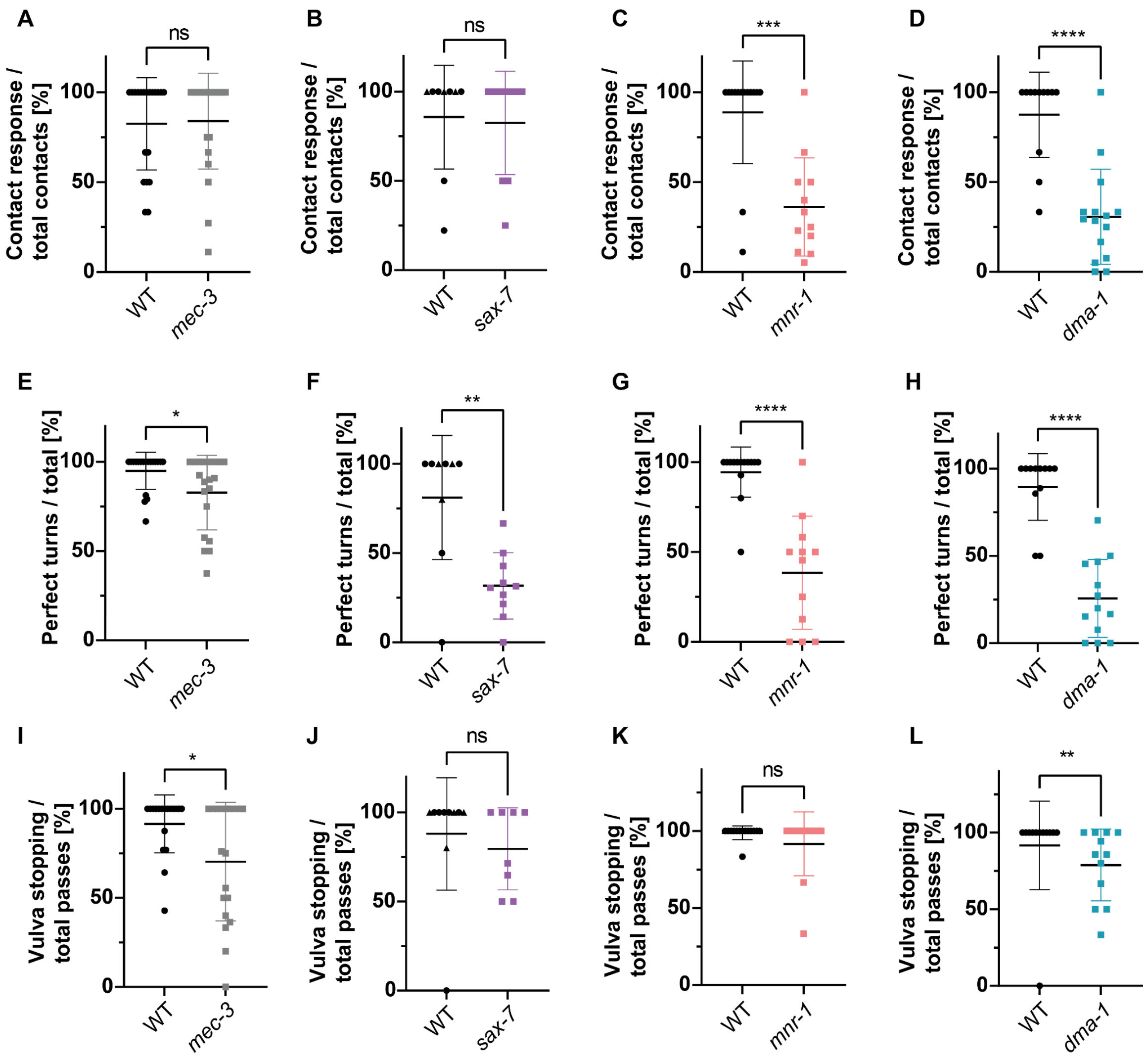

**Fig. S11. *mec-3*, *sax-7, mnr-1* and *dma-1* mutants show male mating behavior defects.** (A,E,I) Hermaphrodite recognition, turning and vulva recognition, respectively, in *mec-3(e1338); him-5(e1490); ser2prom3*::GFP (n = 21, 20,19, respectively) vs WT *him-5(e1490); ser2prom3*::GFP animals (n = 20,19,18, respectively).

(B,F,J) Hermaphrodite recognition, turning and vulva recognition, respectively, in *sax-7(dz156); pF49H12.4*::GFP (n = 10, 10, 8, respectively) vs WT *him-5(e1490); ser2prom3*::GFP and *him-5(e1490); ser2prom3::kaede* (triangle) animals (n = 6+3,6+3,6+4 respectively).

(C,G,K) Hermaphrodite recognition, turning and vulva recognition, respectively, in *mnr-1(dz175); pF49H12.4*::GFP (n = 12 in all panels) vs WT *him-5(e1490); ser2prom3*::GFP animals (n =14 in all panels).

(D,H,L) Hermaphrodite recognition, turning and vulva recognition, respectively, in *dma-1(tm5159); pF49H12.4*::GFP (n = 15,13,12 respectively) vs WT *him-5(e1490); ser2prom3*::GFP (n =12 in all panels). Perfect turn is defined as a turn completed successfully without the male tail losing contact with the hermaphrodite. Mann-Whitney test, black bars show mean, error bars show ± SD. $P$ <0.05 (*), $P$ <0.01 (**), $P$ <0.001 (***); $P$ <0.0001 (****).

**Table S1. List of strains used in this study**

| Name | Genotype | Source |
|---|---|---|
| **N2** | *C. elegans* WT N2 | ***Caenorhabditis* Genetics Center (CGC)** |
| **BP709** | *hmnIs133[ser2prom3::Kaede]* | **Kravtsov et al., 2017** (26)**, Yip and Heiman, 2016** (27) |
| **BP1021** | *hmnIs133[ser2prom3::Kaede]; him-5(e1490) V* | **Inberg et al., 2024** (28) |
| **BP75** | *eff-1(hy21) II* | **Oren-Suissa et al., 2010** (20) |
| **BP462** | *eff-1(ok1021) II; mec-4(e1611) X* | **Podbilewicz Lab** |
| **DR466** | *him-5(e1490) V* | **CGC** |
| **EM253** | *mab-20(bx61ts) I; him-5(e1490) V* | **CGC**<br>**Baird et al., 1991** (29) |
| **EM128** | *mab-21(bx53) III; him-5(e1490) V* | **CGC**<br>**Baird et al., 1991** (29) |
| **NC1687** | *wdIs52[pF49H12.4::GFP unc-119(+)]* | **CGC**<br>**Smith et al., 2010** (30) |
| **EB1564** | *dma-1(tm5159) I; wdIs52[pF49H12.4::GFP unc-119(+)] II;* | **Y. Salzberg**<br>**Salzberg et al., 2013** (31) |
| **EB1653** | *wdIs52[pF49H12.4::GFP unc-119(+)] II; sax-7(dz156) IV* | **Y. Salzberg**<br>**Salzberg et al., 2013** (31) |
| **EB1271** | *wdIs52[pF49H12.4::GFP unc-119(+)] II; mnr-1(dz175) V* | **Y. Salzberg**<br>**Salzberg et al., 2013** (31) |
| **EB1982** | *dzIs53[pF49H12.4::mCherry] II* | **Y. Salzberg**<br>**Ramirez et al., 2019** (32) |
| **MOS81** | *bxIs14[pkd-2::GFP pha-1(+)] him-5(e1490) V* | **M. Oren**<br>**Lints et al., 2004** (9)**, Barr and Sternberg 1999** (10) |
| **MF288** | *ser2prom3::deg-3(u662)* (integrated) | **M. Treinin**<br>**Albeg et al., 2011** (16) |
| **TV15911** | *wyIs592[ser2prom3::myr-GFP odr-1::RFP] III* | **K. Shen**<br>**Dong et al., 2016** (33) |
| **TV15916** | *wyIs581[ser2prom3::myr-mCherry odr-1::GFP] IV* | **K. Shen**<br>**Dong et al., 2016** (33) |
| **TV19321** | *lect-2(ok2617) II; wyIs592[ser2prom3::myr-GFP odr-1::RFP] III* | **K. Shen**<br>**Zou et al., 2016** (34) |
| **TH502** | *ddIs290[sax-7::TY1::EGFP::3xFLAG*(92C12) *unc-119(+)]* | **CGC**<br>**Tang et al., 2021** (35) |
| **CB1338** | *mec-3(e1338) IV* | **CGC** |
| **DA509** | *unc-31(e928) IV* | **M. Oren**<br>**Oren-Suissa et al., 2016** (36) |
| **MT5383** | *lin-44(n1792) I* | **M. Oren** |

| | | |
|---|---|---|
| | | **Miller and Portman, 2011** (37) |
| **PT2565** | *IqIs3(osm-6p::GFP) IV; him-5(e1490) V* | **M. Oren**<br>**Barrios et al., 2012** (38) |
| **PS3380** | *mnIs17(osm-6p::OSM-6::GFP); him-5(e1490) V* | **A. Barrios**<br>**Wang et al., 2010** (23) |
| **EM1106** | *bxIs22[pkd-2::ICE unc-122::GFP]; bxIs14[pkd-2::GFP pha-1(+)] him-5(e1467) V* | **A. Barrios**<br>**Barrios et al., 2008** (39) |
| **BP2283** | *ser2prom3::deg-2(u662)* (integrated); *him-5(e1490); hmnIs133[ser2prom3::Kaede]* | This work<br>MF288 x BP1021♂ |
| **BP2285** | *wyIs592[ser2prom3::myr-GFP podr-1::RFP] III; him-5(e1490) V* | This work<br>TV15911 x (N2 x BP1021 ♂) ♂ |
| **BP2287** | *dzIs53[pF49H12.4::mCherry] II; bxIs14[pkd-2::GFP pha-1(+)]; him-5(e1490) V* | This work<br>EB1982 x MOS81♂ |
| **BP2290** | *wyIs592[ser2prom3::GFP podr-1::RFP] III; mec-3(e1338) IV; him-5 (e1490) V* | This work<br>BP2279 x BP2285 ♂ |
| **BP2291** | *eff-1(ok1021) II; wyIs592[ser2prom3::myr-GFP podr-1::*RFP*] III; him-5 (e1490) V;* | This work<br>BP462 x BP2285 ♂ |
| **BP2293** | *mab-20(bx61) I*; *wyIs592[ser2prom3::myr-GFP podr-1::RFP] III; him-5(e1490) V* | This work<br>EM253 x BP2285 ♂ |
| **BP2294** | *ser2prom3::deg-3(u662)*(integrated); *bxIs14[pkd-2::GFP pha-1(+)] him-5(e1490) V; hmnIs133[ser2prom3::Kaede]* | This work<br>BP2283 x MOS81 ♂ |
| **BP2295** | *eff-1(hy21) II*; *wyIs592[ser2prom3::myr-GFP podr-1::RFP] III; him-5(e1490) V* | This work<br>BP75 x BP2285 ♂ |
| **BP2296** | *eff-1(hy21) II*; *bxIs14[pkd-2::GFP pha-1(+)] him-5(e1490) V* | This work<br>BP2295 x MOS81 ♂ |
| **BP2297** | *ddIs290[sax-7::TY1::EGFP::3xFLAG(92C12) + unc-119(+)]; him-5(e1490) V* | This work<br>TH502 x DR466 ♂ |
| **BP2298** | *dzIs53[pF49H124::mCherry]/+ II; ddIs290[sax-7::TY1::EGFP::3xFLAG(92C12) + unc-119(+)] /+; him-5(e1490) V* | This work<br>EB1982 x BP2297 ♂ |
| **BP2299** | *wyIs592[ser2prom3::myr-GFP; podr-1::RFP] III; him-5(e1490) V; hyEx415[pYI8(ser2prom3::tra-2(ic)::SL2::2xNLS-TagRFP) 20 ng/ul pmyo-2::GFP 10 ng/ul pBluescript KS 70 ng/ul]* | This work<br>BP2285 microinjection |
| **BP2532** | *wyIs592[ser2prom3::myr-GFP; podr-1::RFP] III; him-5(e1490) V; hyEx442[pYI8(ser2prom3::tra-2(ic)::SL2::2xNLS-TagRFP) 20 ng/ul pmyo-2::GFP 10 ng/ul pBluescript KS 70 ng/ul]* | This work<br>BP2285 microinjection |

| | | |
|---|---|---|
| **BP2355** | *wdIs52[pF49H12.4::GFP unc-119(+)] II; sax-7(dz156) IV ;him-5(e1490) V* | This work<br>EB1653 x DR466 ♂ |
| **BP2356** | *wdIs52[pF49H12.4::GFP unc-119(+)] II; sax-7(dz156) IV ;him-5(e1490) V; hyEx445: [dpy-7p::sax-7(s) 20 ng/ul myo-2p::GFP 10 ng/ul pBluescript KS 70 ng/ul] l* | This work<br>BP2355<br>microinjection |
| **BP2356** | *wdIs52[pF49H12.4::GFP unc-119(+)] II; sax-7(dz156) IV ;him-5(e1490) V; hyEx445 [dpy-7p::sax-7(s) 20 ng/ul myo-2p::GFP 10 ng/ul pBluescript KS 70 ng/ul] l* | This work<br>BP2355<br>microinjection |
| **BP2364** | *lin-44(n1792) I; wyIs592[ser2prom3::myr-GFP podr-1::RFP] III; him-5(e1490) V* | This work<br>MT5383 x BP2285 ♂ |
| **BP2366** | *dzIs53[pF49H12.4::mCherry] II; mab-21(bx53) III; bxIs14[pkd-2::GFP pha-1(+)] him-5(e1490) V* | This work<br>EM128 x BP2287 ♂ |
| **BP2369** | *dzIs53[pF49H12.4::mCherry] II; IqIs3(osm-6p::GFP) IV; him-5(e1490) V* | This work<br>EB1982 x PT2565 ♂ |
| **BP2370** | *wyIs581[ser2prom3::myr-mCherry; podr-1::GFP] IqIs3(osm-6p::GFP) IV; him-5(e1490) V* | This work<br>TV15916 x PT2565 ♂ |
| **BP2371** | *bxIs22[pkd-2::ICE unc-122::GFP]; dzIs53[pF49H12.4::mCherry] II; bxIs14[pkd-2::GFP pha-1(+)] him-5(e1467) V* | This work<br>EM1106 x BP2287 ♂ |
| **BP2372** | *mnIs17(osm-6p::osm-6::GFP); dzIs53[pF49H12.4::mCherry] II; him-5(e1490) V* | This work<br>EB1982 x PS3380 ♂ |

**Table S2. List of DNA constructs used in this study**

| Name | Structure | Source |
|---|---|---|
| **pYI08** | *ser2prom3::tra-2(ic)::SL2::2xNLS-TagRFP* | This paper<br><br>pMO32 from Oren-Suissa et al., (36) |
| **pXD22** | *dpy-7p::sax-7(s)* | Dong *et al.,* (40) |

**Table S3.** ***eff-1*** **approximate development time in hours inferred from WT development ratios** (19)

| *eff-1* **hours from egg laid** | **15°C (Values by 16°C)** | **20°C** | **25°C** | **Room temperature 22-23°C** |
|---|---|---|---|---|
| **L4/Adult molt** | 114 | 72 | 55 | 63 |
| **Mid L4** | 99 | 63 | 48 | 55 |
| **L4 + 24 h (1 day adult)** | 123 | 87 | 72 | 78 |

**Table S4. RNA expression levels of characteristic ciliated-neuron genes**

| | | Data from (41) | Data from CeNGEN (15, 42–45) | | | |
|---|---|---|---|---|---|---|
| **Genes selected from** (46) | **Gene description** | **PVD+FLP data Male/Hermaphrodite** | **PVD** | **FLP** | **PDE** | **PHA** |
| ***klp-11*** | Kinesin-II (heterotrimeric) | 0/0 | 78.89 | 0.00 | 9.06 | 31.79 |
| ***kap-1*** | Heterotrimeric kinesin-II associated protein | 0.015/0.00 | 181.06 | 137.63 | 107.26 | 163.57 |
| ***daf-19*** | RFX transcription factor | 0.005/0.02 | 21.22 | 61.70 | 141.11 | 36.17 |
| ***bbs-2*** | BBS2 | 0/0 | 0.00 | 0.00 | 482.62 | 255.21 |
| ***klp-20*** | Kinesin-II (heterotrimeric) | 0/0 | 19.02 | 0.00 | 0.00 | 22.25 |
| ***bbs-5*** | BBS5 | 0/0 | 0.00 | 0.00 | 508.17 | 379.25 |
| ***che-2*** | IFT80 | 0/0 | 0.00 | 0.00 | 126.81 | 107.92 |
| ***osm-5*** | IFT88 | 0/0 | 0.00 | 0.00 | 1138.27 | 621.59 |
| ***ift-81*** | IFT81 | 0/0 | 0.00 | 0.00 | 1629.16 | 593.09 |
| ***osm-1*** | IFT172 | 0/0 | 3.49 | 12.99 | 291.95 | 52.94 |
| ***che-3*** | IFT-dynein | 0/0 | 0.00 | 0.00 | 148.12 | 210.13 |
| ***che-13*** | IFT57 | 0/0 | 0.00 | 0.00 | 563.12 | 426.45 |
| ***bbs-1*** | BBS1 | 0/0 | 0.00 | 0.00 | 34.41 | 170.92 |
| ***bbs-8*** | BBS8 | 0/0 | 0.00 | 0.00 | 349.32 | 478.58 |
| ***osm-6*** | IFT52 | 0/0 | 0.00 | 0.00 | 345.16 | 358.82 |
| ***che-11*** | IFT140 | 0/0 | 0.00 | 0.00 | 491.14 | 133.45 |
| ***xbx-1*** | Dynein light intermediate chain | 0/0 | 0.00 | 0.00 | 227.05 | 148.83 |
| ***osm-9*** | Olfactory channel | 0/0 | 19.02 | 0.00 | 0.00 | 15.86 |
| ***osm-3*** | Kinesin-II (homodimeric) | 0/0 | 0.00 | 0.00 | 8.40 | 22.47 |
| ***tax-2*** | Cyclic nucleotide gated olfactory channel | 0/0 | 0.00 | 0.00 | 0.00 | 0.00 |

**Movie legends**

**Movie S1. Hermaphrodite traced PVD neuron rotation.**
Three dimensional unilateral reconstruction of PVD branching orders in the cell body region of a three-day old hermaphrodite *him-5(e1490); ser2prom3::GFP* animal. Gray: axon; blue: primary (1°) branch; purple: secondary (2°) branches; red: tertiary (3°) branches; cyan: quaternary (4°) branches (which extend laterally) and ectopic quaternary branches (which extend medially); pink: ectopic secondary branches, which do not reach the tertiary line; light pink: ectopic tertiary branches, which branch from an ectopic secondary process; yellow: ectopic fifth-order (5°) branches, which branch from a quaternary. Worm width at the axon region is 85 μm and visible primary length is 225 μm.

**Movie S2. Male traced PVD neuron rotation.**
Three dimensional unilateral reconstruction of PVD branching orders in the tail region of a three-day old male *him-5(e1490); ser2prom3::GFP* animal. Blue: primary (1°) branch and axon; purple: secondary (2°) branches; red: tertiary (3°) branches; cyan: quaternary (4°) branches (which extend laterally) and ectopic quaternary branches (which extend medially); pink: ectopic secondary branches, which do not reach the tertiary line; light pink: ectopic tertiary branches, which branch from an ectopic secondary process; yellow: ectopic fifth-order branches (5°), which branch from a quaternary. Worm width at the axon region is 47 μm and visible primary length is 205 μm.

**Movie S3. Male traced PVD and RnB in the L4.**
Three dimensional reconstruction of a male L4 *dzIs53[pF49H12.4::mCherry]*; *bxIs14[pkd-2::GFP]*; *him-5(e1490)* animal. PVD is traced in magenta while RnB (except R6B, which is not marked by *pkd-2::GFP* (8) is traced in green. Worm width is 35 μm, visible primary length is 97 μm.

**Movie S4. PVD and RnB do not colocalize.**
Three-dimensional image Z-series of the animal shown in Figure 2 *B*. Ventral view of an adult WT 1-2 day-old *him-5(e1490)* male expressing PVD (*pF49H12.4::mCherry*) and RnB (*pkd-2::GFP*) markers. Scale bar is 25 μm, Z-slices are spaced 0.6 μm apart.

**Movie S5. Mating of WT young adult to adult transition.**
Side-by-side time-synchronized representative mating segments of a young adult male *ser2prom3::GFP*; *him-5(e1490)* animal (left) and the same animal 24 h later (right), mating with *unc-31(e927)* hermaphrodites (variable magnification, hermaphrodites are ~1mm in length).

**Movie S6. PVD growth into the male tail rays.**
Extended timelapse images of the animal shown in Figure S4. Timelapse in minutes highlighting two growth events of ectopic processes in ray 6 (orange and purple arrowheads). Animal is a young adult WT (a few hours after molting) *ser2prom3::GFP; him-5(e1490)* male mounted in 0.05% tetramisole on a 5% agarose pad. Signal is inverted. Scale bar is 25 μm.
